# Host-derived Interleukin 1α induces an immunosuppressive tumor microenvironment via regulating monocyte-to-macrophage differentiation

**DOI:** 10.1101/2024.05.03.592354

**Authors:** Manikanda Raja Keerthi Raja, Gourab Gupta, Grace Atkinson, Katie Kathrein, Alissa Armstrong, Michael Gower, Igor Roninson, Eugenia Broude, Menqiang Chen, Hao Ji, Chang-uk Lim, Hongjun Wang, Daping Fan, Peisheng Xu, Jie Li, Gang Zhou, Hexin Chen

## Abstract

Tumor-associated macrophages exhibit high heterogeneity and contribute to the establishment of an immunosuppressive tumor microenvironment (TME). Although numerous studies have demonstrated that extracellular factors promote macrophage proliferation and polarization, the regulatory mechanisms governing the differentiation process to generate phenotypically, and functionally diverse macrophage subpopulations remain largely unexplored. In this study, we examined the influence of interleukin 1α (IL-1α) on the development of an immunosuppressive TME using orthotopic transplantation murine models of breast cancer. Deletion of host Il1α led to the rejection of inoculated congenic tumors. Single-cell sequencing analysis revealed that CX3CR1+ macrophage cells were the primary sources of IL-1α in the TME. The absence of IL-1α reprogrammed the monocyte-to-macrophage differentiation process within the TME, characterized by a notable decrease in the subset of CX3CR+ ductal-like macrophages and an increase in iNOS-expressing inflammatory cells. Comparative analysis of gene signatures in both human and mouse macrophage subsets suggested that IL-1α deficiency shifted the macrophage polarization from M2 to M1 phenotypes, leading to enhanced cytotoxic T lymphocyte activity in the TME. Importantly, elevated levels of IL-1α in human cancers were associated with worse prognosis following immunotherapy. These findings underscore the pivotal role of IL-1α in shaping an immune-suppressive TME through the regulation of macrophage differentiation and activity, highlighting IL-1α as a potential target for breast cancer treatment.

**Teaser:** Interleukin 1α dictates macrophage behavior, influencing an immunosuppressive microenvironment in breast cancer, suggesting it as a treatment target.

## Introduction

Immunotherapy is rapidly evolving as a treatment approach for breast cancer, however, the response rates achieved thus far have been suboptimal(*1*). It is widely recognized that the immunosuppressive nature of the tumor microenvironment (TME) plays a crucial role in limiting the efficacy of therapies targeting breast tumors(*2–4*). Tumor growth can trigger an influx of bone marrow-derived monocytes, which differentiate into monocytic myeloid-derived suppressor cells (Mo-MDSCs), monocyte-derived dendritic cells (Mo-DCs), and tumor-associated macrophages (TAMs)(*5, 6*). These cells contribute to creating an immunosuppressive tumor microenvironment (TME). Monocyte-macrophage lineage cells display a high degree of plasticity and heterogeneity(*7–9*), and meanwhile they share expression of numerous surface markers and common properties; therefore, it is challenging to clearly distinguish and characterize these cell populations(*10*).

TAMs are the most abundant immune cells in TME of many types of tumors including breast cancer and can be roughly classified as M1 and M2 macrophages(*11, 12*). M1 macrophages play a crucial role in antitumor immunity and primarily mediate proinflammatory processes, whereas M2 macrophages have been demonstrated to have protumor features(*13*). However, TAMs may exhibit a wide range of differentiation and activation states, with the classical M1 and M2 phenotypes representing the extreme ends of this spectrum(*14*). *In vitro* studies have revealed that factors such as M-CSF, LPS, IFN-γ, and IL-4 can further polarize macrophages into additional M1 or M2-like subsets(*14*), likely reflecting the circumstances of *in vivo* differentiation. Recent single cell sequencing analyses have identified many more subsets of macrophage *in vivo*(*15–18*). Interestingly, these macrophage subsets identified by single cell sequencing analysis generally do not reflect “M1/M2” polarization(*16, 17*), most likely due to exposure to multiple spatial and temporal stimuli and cellular context. Furthermore, it also remains largely unknown how intrinsic factors regulate monocytes/macrophages for the fate determination and functional adaptation in response to the dynamic extrinsic microenvironmental cues(*8*).

Interleukin-1s (IL-1s) as the potent apical cytokines instigate multiple downstream processes to affect both innate and adaptive immunity. Both IL-1α and IL-1β bind to the same receptor, a type 1 IL-1 receptor (IL-1R), to activate the downstream signaling cascade (*19*). In contrast to the highly restricted expression of IL-1β in immune cells, IL-1α is constitutively expressed in epithelial, endothelial, and stromal cells (*19*). Notably, tumor-derived IL-1α can affect cancer progression by acting on both tumor and immune cells(*20*). Our recent work revealed an essential role of IL-1α in tumorigenesis in producing a chronic inflammatory environment conducive to the maintenance of cancer stem cells(*21*). Kuan et al reported that breast tumor-derived IL-1α acts on tumor-infiltrating myeloid cells to induce the expression of thymic stromal lymphopoietin (TSLP), which in turn promotes the survival of tumor cells(*22*). Secreted IL-1α can promote the production of neutrophils and less mature CD11b+ myeloid cells resulting in systemic and local immune suppressive environment(*2*). In contrast to tumor-promoting function in a variety of cancers, a few have found that particularly transient expression of membrane-bound IL-1α in fibrosarcoma and lymphoma cells might exhibit anti-tumorigenic effects(*23–26*). As a matter of fact, very limited studies have been conducted to reveal the functions of host-derived IL-1α in tumorigenesis(*27–29*). Intriguingly, whole-body knockout of IL-1α has the opposite effects on oncogene Her2/neu and polyoma T antigen-induced mammary tumor in transgenic mouse models(*21, 30*). It is most likely that different sources of IL-1α mediated dynamic immune responses during tumorigenesis.

In our current investigation, utilizing congenic transplantation models, we examined the role of host-derived-IL-1α in breast tumorigenesis via reprogramming immunosuppressive myeloid cells. Using both single cell sequencing and flow cytometry analysis, we uncovered the heterogeneity of the monocyte and macrophage-subpopulations and elucidated the mechanisms underlying the rejection of tumor challenge in the Il-1α deficient mice. Gaining insights into this common immunosuppressive mechanism mediated by host-derived IL-1α may pave the way for the development of novel immunotherapeutic strategies against breast cancer.

## Results

### Il1α-deficient mice hindered the growth of transplanted tumors and exhibited modified local and systemic immune responses

Our previous study has shown that knockout of IL1α inhibit MMTV-neu induced tumorigenesis(*21*). To assess the impact of host-derived IL1α on tumor progression, we performed orthotopic injections of H605 cells derived from a MMTV-neu tumor into the mammary pads of both wild type (WT) and *Il1α^-/-^* mice. Tumor growth was monitored at three-day intervals over a 40-day period. Intriguingly, the majority of transplanted tumors in *Il1α ^-/-^* mice grew for approximately two weeks followed by regression, whereas those in the WT mice continued to grow (Fig. 1A).

**Fig. 1.**
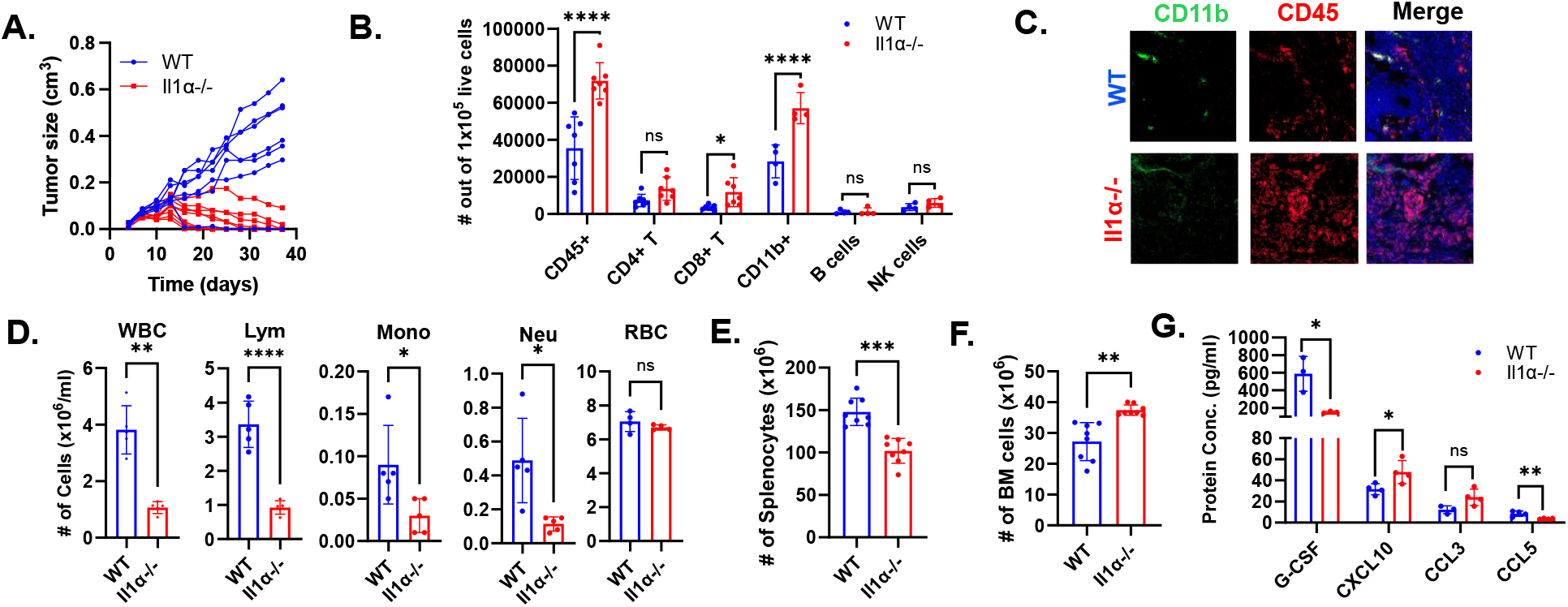
Il1α absence affects mammary gland tumor growth and systemic immunity. **A.** H605 tumor growth in WT and *Il1α^-/-^* (n = 7). **B.** Flow cytometry analysis of immune cells in H605 transplanted tumors at 2 weeks after inoculation. The number of infiltrating immune cells was calculated from 100,000 live cells using flow cytometry (n = 7). **C.** Representative immunofluorescence staining of CD11b and CD45 markers from WT and *Il1α^-/-^* tumor sections. **D**. Blood Vetscan® data shown between tumor-bearing WT and *Il1α^-/-^* mice. Unpaired Two Tailed t-test was performed. **<0.01; *<0.05. **E** and **F** describe the total number of spleen and bone marrow leukocytes from tumor-bearing WT and *Il1α^-/-^* mice. **G.** Concentration of cytokines GCSF, CXCL10, CCL3 and CCL5 in the sera of tumor-bearing WT and *Il1α^-/-^* mice. Unpaired Two Tailed t-test was performed. **<0.01; *<0.05.

To better elucidate the underlying mechanisms, we focused on the HER2-positive breast cancer model for further investigation and specifically studied the tumor microenvironment at the two-week time point when the tumor regression became apparent. Dissected tumors were digested into single cells and subjected to flow cytometry to characterize the infiltrated immune cells (Fig. S1). Remarkably, *Il1α^-/-^* tumors exhibited a significant increase in CD45+ tumor-infiltrating leukocytes (Fig. 1B-C). Within those leukocytes, both CD8+ T cells and CD11b+ myeloid cells were notably elevated in tumors from *Il1α^-/-^*mice compared to WT mice.

Considering that IL-1α plays a role in regulating hematopoiesis and the margination of immune cells(*31, 32*), we investigated the systemic changes in the immune systems. Interestingly, both tumor-free and tumor-bearing *Il1α^-/-^*mice showed a significant decrease in overall blood leukocyte output compared to corresponding WT mice (Fig.1D and Fig. S2). The skewed blood output revealed an increased presence of B cells and Ly6C+ monocytes, accompanied by a reduction in CD4, CD8, and neutrophils (Fig. S3A-B), which differed from the distribution of infiltrated immune cells in the tumor. We reasoned that the altered periphery immune cell distribution might stem from abnormal hematopoiesis in the spleen and bone marrow(*31*). Indeed, in the tumor-bearing *Il1α^-/-^* mice, spleen cellularity was diminished compared to WT mice. The lymphoid distribution mirrored the blood profile trend, while the myeloid cell numbers remained unchanged (Fig. 1E and Fig. S4A-B). Conversely, in tumor-bearing *Il1α^-/-^* mice, the bone marrow exhibited increased cellularity, featuring more B cells and fewer neutrophils, consistent with the patterns observed in the spleen and blood (Fig. 1F and Fig. S5A-B).

To gain a deeper understanding of the altered immune cell distribution among organs, we conducted ELISA analysis of blood plasma, revealing barely detectable IL1α in WT mice (Fig. S6). There was a decreasing trend in cytokines including granulocyte-colony stimulating factor (G-CSF), chemokine C-C ligand 5 (CCL5), and increased expression of C-X-C motif chemokine ligand 10 (CXCL10) (Fig. 1G). Reduced levels of G-CSF and CCL5 may contribute to the observed leukopenia by limiting egression of immune cells from bone marrow to periphery systems(*32–34*). Additionally, cytokines like CXCL10 are required for T cell recruitment to local site of inflammation(*35*), partially explaining the abundant recruitment of T cells to the *Il1α^-/-^* TME. Collectively, these results indicated that Il1α-deficient mice displayed altered local and systemic immune responses against tumor development.

### Loss of IL1α leads to an immune-active TME

The intricate interplay of immune, cancer, and stromal cells defines the complexity of the TME. To explore the influence of IL-1α on the TME, we conducted single-cell RNA sequencing (scRNA-seq) analysis on two-week tumor samples from both WT and *Il1α^-/-^* mice, unveiling the existence of 14 distinct clusters (Fig. 2A and Fig. S7). Through a UMAP representation comparison, we observed a significant increase in the CD8 cluster and a decrease in the tumor cluster in *Il1α^-/-^*mice. Further analysis of differentially expressed soluble factors and their receptor levels revealed striking disparities between the WT and *Il1α^-/-^* conditions. Among nonimmune cell clusters, the tumor cluster in *Il1α^-/-^* mice showed a remarkable upregulation of chemokines, specifically Cxcl9 and Cxcl10, with the expression levels reaching up to 75%, whereas they were barely detectable in WT (Fig. 2B). Additionally, nonimmune clusters like endothelial cells (Endo) and fibroblasts (Fibro) in the *Il1α^-/-^* mice exhibited lower expression levels of interleukin 33 (*Il-33*) and colony-stimulating factor 1 (*CSF1*) compared to the WT group. In the immune cell populations, interleukin 16 (*Il-16*)consistently exhibited lower expression in *Il1α^-/-^* mice. Conversely, *Ccl5* and C-X-C motif chemokine receptor 4 (*Cxcr4*) were expressed at higher levels in Other T cells, CD8 cells, NK, and NKT cells from *Il1α^-/-^* mice. Markers associated with T cell activation, such as *CD28*, *Icos*, *Ctla4*, and *Pdcd1*, were significantly upregulated in Other T cells, CD8 cells, NK, and NKT cells in the *Il1α^-/-^* condition. Furthermore, critical T cell cytokines with anti-tumor properties, including interferon gamma (*IFN-γ*), granzyme B (*GzmB*), and tumor necrosis factor (*TNFα*), were significantly expressed in the same lymphoid cells that displayed the activation markers (Fig. 2B). This suggests a potential link between the upregulation of T cell activation markers and the production of anti-tumor cytokines within the *Il1α ^-/-^* TME. Within the myeloid clusters, there was a notable trend of reduced expression of colony stimulating factor receptors, including *Csf1r*, *Csf2ra*, and *Csf3r*, especially in the Mono/Mac cluster. Conversely, the Mono/Mac cluster exhibited higher expression levels of *Cxcr4*, while all myeloid clusters displayed decreased levels of *Tgfb1* in the *Il1α^-/-^* mice (Fig. 2B).

**Fig. 2.**
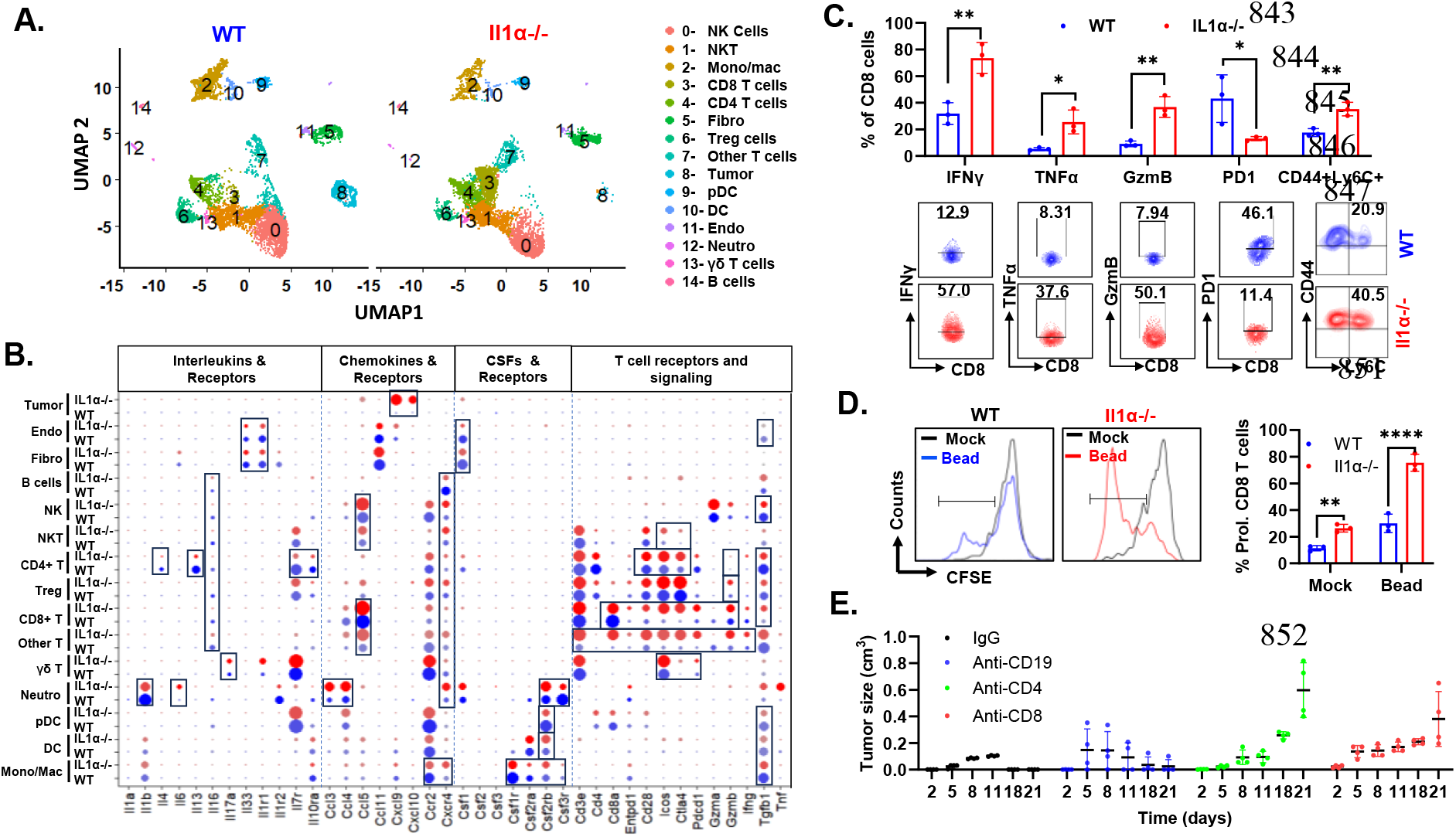
Loss of IL1α results in an immune-active TME. **A.** t-SNE scRNA-seq plot comparing intratumoral cells between WT and *Il1α^-/-^* samples. **B.** Differential expressed soluble factors, and their receptor gene levels found across all clusters shown in dot plot. **C.** Quantification of T cell cytokines, exhaustion marker PD1 and stem like T cell marker in infiltrated CD8 T cells using flow cytometry (n = 3). **D**. Quantification of *ex vivo* tumor T cells proliferation using anti-CD3/CD28 beads (n = 3). Unpaired Two Tailed t-test was performed.**,p<0.01; ****<0.0001. **E.** H605 tumor growth in the transplanted *Il1α^-/-^*mice post depletion of CD19, CD4 and CD8 T cells (n = 4).

To corroborate the scRNA-seq findings and confirm whether T cells were indeed more active and less inhibited in the transplanted tumors of *Il1α^-/-^*mice, we conducted flow cytometry analysis to quantify the pro-tumorigenic cytokine profile and the *ex vivo* anti-CD3/CD28 bead proliferation capacity of T cells isolated from tumors. The flow cytometry results demonstrated a significant elevation in the secretion of potent cytotoxic CD8 T cell cytokines, such as TNFα, IFNγ, and GzmB. Moreover, there was an upregulation in the levels of active T cell indicators like CD44 and Ly6C, alongside a reduction in the expression of exhaustion marker PD-1(Fig. 2C). Furthermore, when cultured *ex vivo*, both stimulated and unstimulated CD8 T cells from tumors in *Il1α^-/-^* mice exhibited a significantly elevated proliferation rate, contrasting starkly with those from the WT group (Fig. 2D). To assess the significance of T cells in the observed tumor regression, we depleted CD4 T, CD8 T, and B cells in the *Il1α^-/-^* mice (Fig. S8). Strikingly, CD4 and CD8 T cell depletion resulted in a different tumor growth pattern compared to WT, with a noticeable lack of exponential growth up to day 14, after which tumor growth resumed (Fig. 2E). Together, these results suggest that the TME in *Il1α^-/-^* mice demonstrated a more immune-active phenotype, contributing to tumor clearance compared to that observed in *WT* mice.

### CX3CR1+ macrophages are the major cellular source of IL-1α in TME

Although the expression levels of IL-1α were generally low (Fig. 2B), the major source of IL-1α in the TME was identified to originate from the Mono/Mac cluster (Fig. 3A). Further scRNA-seq data from mouse breast cancer aligned remarkably well with the human breast cancer scRNA-seq dataset showing that only a small fraction of myeloid cells expressed *IL-1α* in TME (Fig. S9A-C and Fig. 3B). Given that monocytes/macrophages are highly heterogeneous in TME and have diverse functions, our investigation delved deeper by sub-clustering the Mono/Mac cluster to attain a finer resolution for the cellular source of *Il1α* (Fig. S10). As shown in Fig. 3C, the UMAP plot reveals a total of seven subclusters designated as Mono and Mac1 through 6. The Mono cluster was characterized by high levels of monocyte markers such as *Ly6c2* and *Ccr2* and absence/low expression of macrophage markers such as *Adgre1*, *Retnla* and *H2-Aa*. The Mac 1-6 clusters are characterized by different combinations of macrophage markers, reflecting their different differentiation statuses and functions (Fig. 3D).

**Fig. 3.**
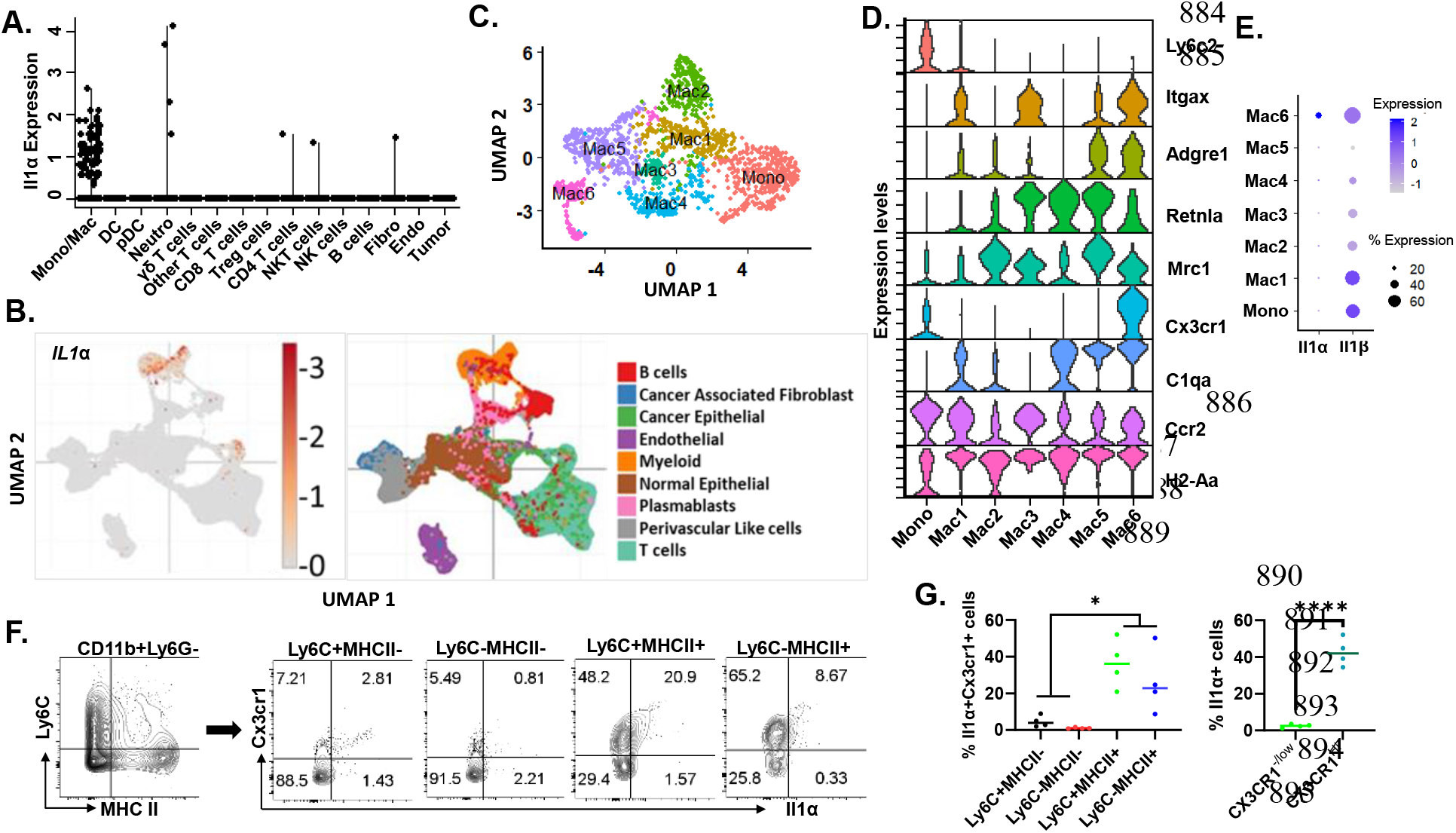
CX3CR1+ macrophages serve as the major cellular source of Il1α in the TME. **A.** Expression of Il1α from TME scRNAseq shown in violin plot. **B.** t-SNE scRNA-seq plot from human breast cancer dataset depicting clusters where *Il1α* production is observed. **C.** t-SNE scRNA-seq plot of reclustered Mono/Mac clusters. **D.** Violin plot illustrating the expression of macrophage markers across the subclusters. **E**. Dot plot illustrating expression levels of Il1α and Il1β in reclustered Mono/Mac clusters (Mac 1 through 6). **F.** Validation of the source of Il1α in the TME through flow cytometry analysis. **G.** Quantification of Il1α-producing myeloid cells from the 2-week transplants in WT mice (n = 4). Unpaired Two Tailed t-test was performed. *<0.05; ****<0.001.

Intriguingly, *Il1β* expression was detected in all clusters and 60% cells in Mono, Mac1, and Mac6 clusters, while factor Il1α was restrictedly expressed in the only 20% cells in Mac 6 cluster (Fig. 3E). Mac6 cluster expressed many macrophage markers including *Adgre1*, *Retnla*, *Mrc1*, *H2-Aa* and *C1qa*. Interestingly, only one marker, *Cx3cr1*, can uniquely identify IL-1α producing macrophages. To validate the scRNA-seq findings and identify the Il1α-producing population, we employed flow cytometry to identify the IL-1α producing cells in the tumors of WT mice. Based on the expression levels of Ly6c and MHC II, the mono/mac cells were stratified into four subpopulations within each quadrant. Only a fraction of MHC II+ cells expressed IL-1α. Consistent with our scRNA-seq data, the majority of IL-1α producing macrophages were CX3CR1^+^MHCII^+^ Ly6C^lo/-^ macrophages (Fig. 3F-G).

### IL-1α skews differentiation of monocytes towards CX3CR1^+^ macrophages

Monocytes recruited into TME can undergo diverse differentiation pathways leading to the development of various macrophage subpopulations(*36, 37*). To investigate the potential regulatory role of IL-1α in this process, analysis on myeloid cell distribution in the TME of tumors from WT and *Il1α^-/-^* mice was conducted. The findings revealed an increased presence of Mac2 and a decrease in Mac4-6 cluster cells in tumor from transplanted *Il1α^-/-^* mice (Fig. 4A-B). Pseudo time trajectory analysis, initiated with the Mono cluster in both *WT* and *Il1α^-/-^* samples, demonstrated a similar trajectory up to Mac1. However, *Il1α^-/-^* mice showed a preference for the Mac2 trajectory, not observed in *WT*, and the Mac3 gave rise to Mac 6 via Mac 4 and Mac5 in *WT*, a progression disrupted in *Il1α^-/-^* ( Fig.4C).

**Fig. 4.**
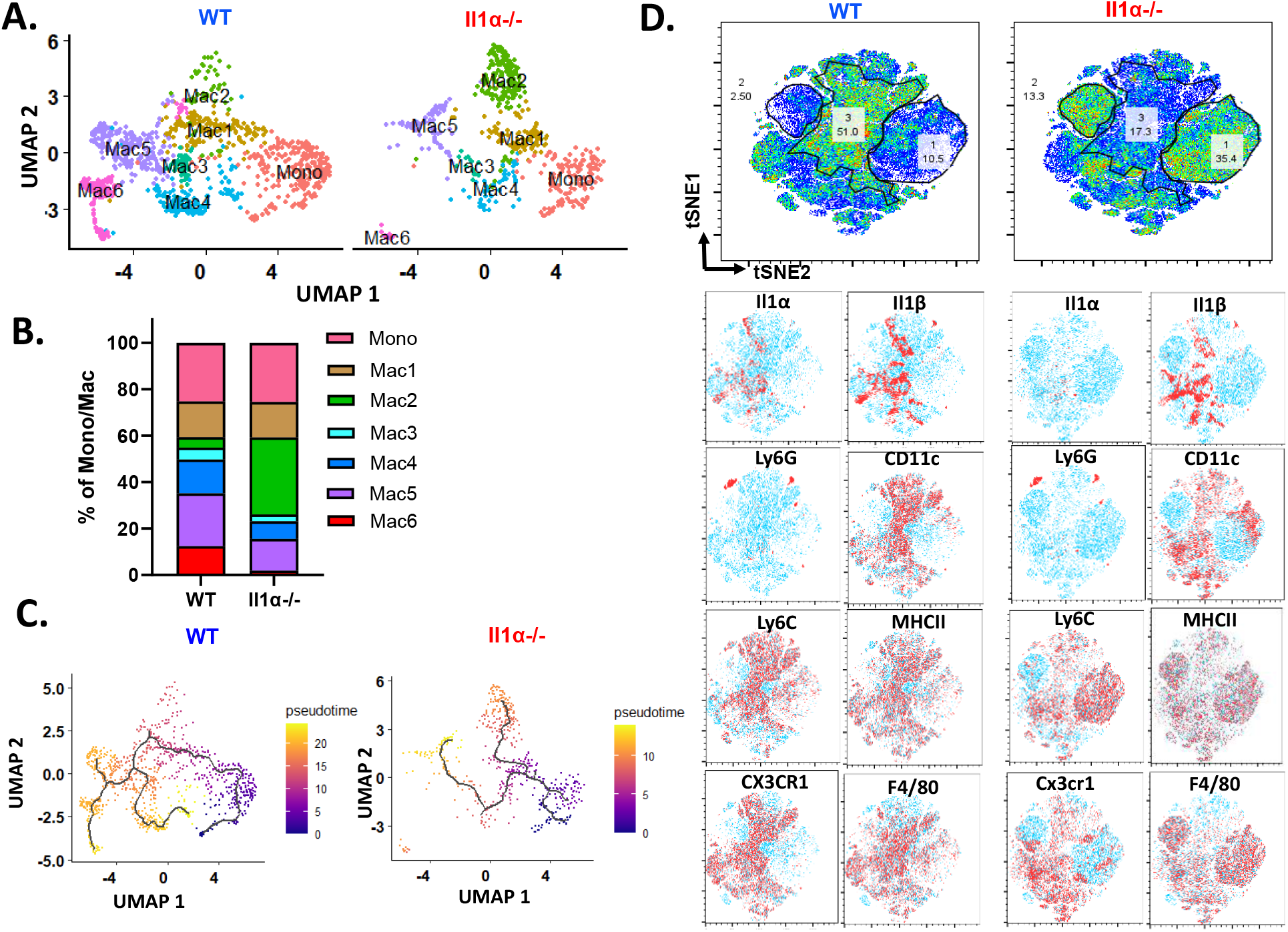
Il1α skews differentiation and alters myeloid phenotype. **A.** t-SNE scRNA-seq plot comparing reclustered Mono/Mac clusters in WT and *Il1α^-/-^*. **B.** Percentage distribution of each subcluster within the total Mono/Mac, comparing WT and *Il1α^-/-^*. **C.** t-SNE scRNA seq trajectory analysis comparing subclusters between WT and *Il1α^-/-^*. **D.** Representative t-SNE visualization from flow cytometry of WT and *Il1α^-/-^*. Gated CD45^+^CD11b^+^ myeloid cells highlighting different markers such as Il1α, Il1β, Ly6G, CD11c, Ly6C, MHCII, CX3CR1 and F4/80.

Mac2 cells expressed the typical macrophage markers including *Adge1* and *Mrc1* but low levels of *Ly6c2* and *H2-Aa* (Fig. 3D), which may represent an intermediate state between monocyte and mature macrophages. In contrast, both Mac5 and Mac6 expressed high levels of *H2-Aa* and *Adge1* and low levels of *Ccr2* and *Ly6c2*, indicative of more mature macrophages. To validate the scRNA-seq results, flow cytometry analysis of monocyte/macrophages in TME was performed. Consistently, the CX3CR1^+^CD11c^+^F4/80^+^ macrophages (population 3), corresponding to Mac6 population, were significantly reduced in tumors from *Il1α^-/-^* mice compared the WT ones (Fig. 4D). Conversely, CX3CR1^-^ macrophage populations (population 1 and 2), including both immature and mature macrophages differing in their Ly6C expression status, increased in transplants from *Il1α^-/-^*mice (Fig. 4D). These results suggested that IL1α influences the differentiation of recruited monocytes toward CX3CR1+ mature macrophages.

### Impacts of IL1α deficiency on the functions of myeloid cells in the TME

Observing the distinct distribution patterns within these Mono/Mac clusters prompted us to conduct Gene Set Enrichment Analysis (GSEA) to gain deeper insights regarding their functions in TME (Fig. 5A). The mono **c**luster exhibited classical monocyte marker genes, such as *Ccr2* and *Ly6c2* (Fig. 5B). Additionally, it shared genes related to pathways involving the response to LPS, regulation of leukocyte cell-cell adhesion, regulation of hemopoiesis, and leukocyte migration (Fig. 5A and Fig. S11A-C). Mac1 population, representing the first branching point in the trajectory (Fig. 4C), showed upregulation of genes associated with pathways like response to LPS, response to type II interferon, and cellular response to IL-1 (Fig. S11A, D-E). This branching point preferentially gave rise to either Mac2 or Mac3. Mac2 had genes upregulated for hypoxia sensing and oxidative stress-responsive genes such as *Vegfa*, *Hilpda*, *Bnip3*, *Egln3*, and *Ndrg1* (Fig. 5B and S11F), a unique feature compared to other subclusters, and expressed genes involved in chemokine response and leukocyte migration (Fig. S11G), implying inflammatory reactions. Despite Mac2 cells exhibiting both M1 and M2 polarization genes, including *Nos2* and *Arg1*, relatively lower levels of mature macrophage markers like *Adge1*, *H2-Aa*, *C1qa* and *Cx3cr1* indicate their immature status (Fig. 5B). Mac3 exhibited upregulated genes such as *St3gal4*, *Tspan32*, *Lilrb4a*, and *Cd24a*, contributing to the regulation of cell-cell adhesion (Fig. S11B). Interestingly, the expression profiles of Mac4 cells did not show enrichment in the analyzed GSEA pathways. Mac5 featured genes like *Trf*, *Tfrc*, *Apoe*, *Ap2a2*, and *Ap2m1*, which are involved in receptor-mediated endocytosis and lymphocyte-mediated immunity (Fig. S11H-I). Mac5 also expressed markers such as *Folr2*, *Mrc1*, *Apoe*, and *C1q* (Fig. 5B), resembling the tissue-resident macrophage(*38, 39*). Mac6 expressed typical monocyte-derived macrophage markers such as *Cx3cr1*, *Trem2*, *Cadm1* and *Spp1* (Fig. 5B), resembling the ductal macrophages known to be involved in apoptotic cell clearance and tissue remodeling, and significantly expanded during tumorigenesis (Fig. S11F)(*38*). The IL1 production pathway was observed most in Mac6 evidently with genes such as *Il1α*, *IL1β* and *Il1rn* (Fig. S11E and S11J).

**Fig. 5.**
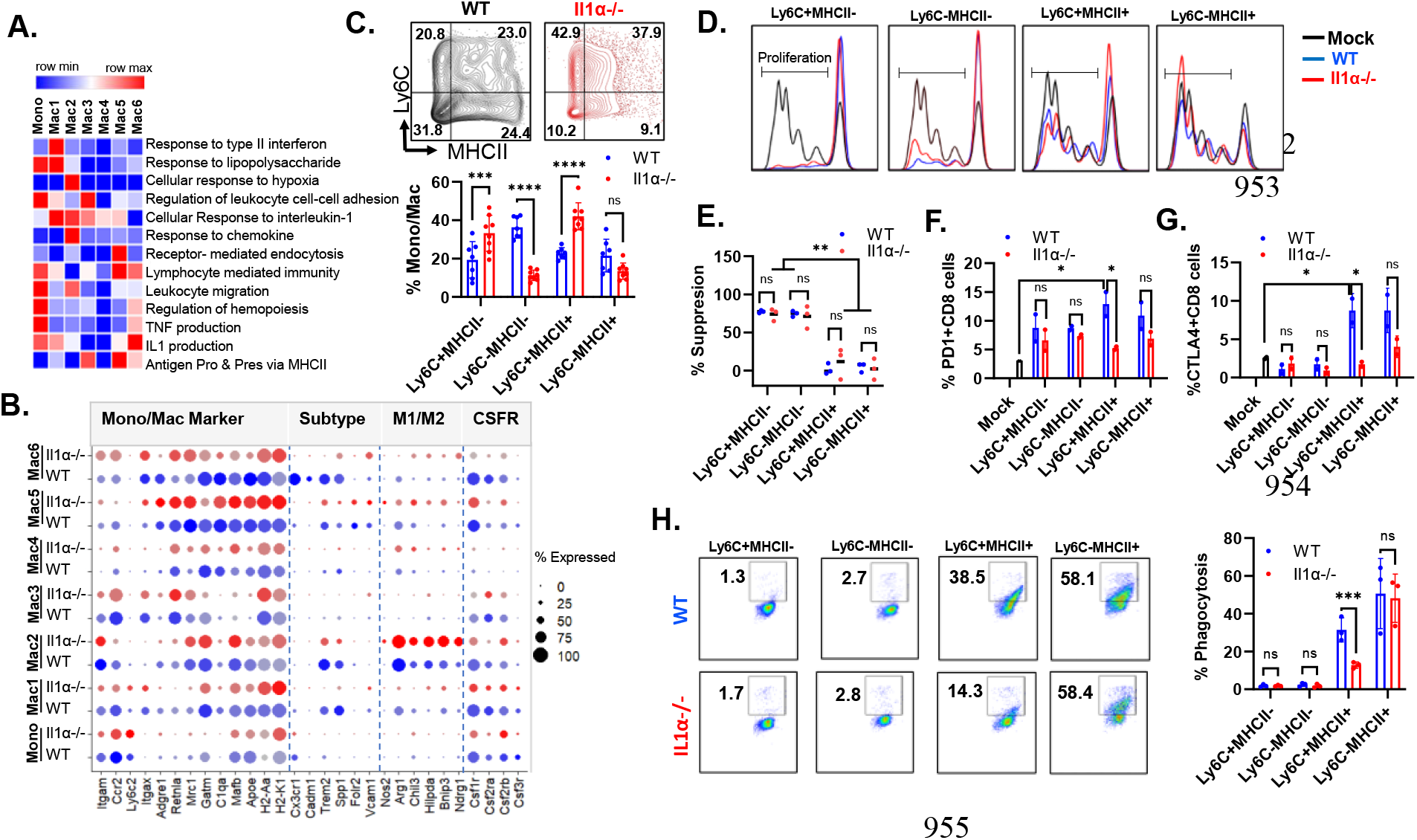
Il1α alters heterogeneity and functions of TAMs. **A.** Heatmap of GO pathway analysis across Mono/Mac subclusters. **B.** scRNAseq Differential expressed genes found in WT and *Il1α^-/-^* myeloid subclusters. **C.** flow cytometry analysis of CD45^+^CD11b^+^Ly6G^-^ gated myeloid cells distribution based on the expression levels of Ly6C and MHCII (n = 5). 2-way ANOVA. ***<0.05; ****<0.001. **D.** Representative figure depicting bead mediated T cell proliferation co-culture with each quadrant of myeloid population shown in Fig. 5C. **E.** Quantification of T cell suppression mediated by each quadrant of myeloid population. 2-way ANOVA. **<0.006. **F** and **G**. Quantification of exhaustion markers PD1 and Ctla4 mediated by myeloid cell on co-cultured T cells (n=2). 2-way ANOVA. *<0.006. **H**. Phagocytosis assay of micro-beads with each quadrant of myeloid population and representative flow cytometry graph (n =3). 2-way ANOVA. ns, not significant; ***<0.001.

In the *Il1α^-/-^* TME, there was an elevation in the Mac2 population, along with the declines in the Mac 4-6 populations. Furthermore, macrophages exhibited decreased expression of key macrophage differentiation markers, including *Adgre1*, *C1qa*, *C1qb*, *C1qc*, *Gatm*, and *Apoe* within Mac1, Mac4, and Mac6 subclusters (Fig. 5B). Furthermore, clusters of Mono, Mac1, Mac2 and Mac6 in *Il1α^-/-^* mice showed diminished expression of receptors essential for differentiation, such as *Csf1r*, *Csf2ra*, and *Csf3r*. Notably, the M1 macrophage marker genes such *Nos2, Bnip3* and *Ndrg1* gene were upregulated especially in clusters of Mac2, Mac5 and Mac6 in *Il1α^-/-^* mice (Fig. 5B).

Given the stark differences in the cell populations and marker gene expression between *WT* and *Il1α^-/-^*, we adopted a common strategy employing the Ly6C *vs*. MHC II gate to study the functional aspects of subpopulations within each quadrant. Ly6c^+^MHC II^-^ cells may contain Ly6C^+^ monocytes. The Ly6C^+^MHCII^+^ cells, characterized by the expression of Ly6C, MHCII, CD11c, and a low level of CX3CR1 (Fig. 4D), may encompass cells in Mac2-3 clusters. Notably, we observed the phenotypical shift on the Ly6C^+^MHCII^+^ cells from MHC II^hi^ to MHC II^lo^, and such a shift might result from the changes of cellular composition. The Ly6C^-^MHCII^-^ cells, exhibiting low levels of Ly6C, MHCII, and CX3CR1, bear the resemblance to Mac4 cells. The Ly6C^-^MHCII^+^ cells may include cells from Mac5 and 6 clusters. Consistent with the increase in Mac2 and reduction in Mac 4-6 in the *Il1α^-/-^* TME, flow cytometry analysis confirmed the reduced Ly6C^-^ MHCII^-^ and Ly6C^-^MHCII^+^ cells but increased Ly6C^+^MHCII^+^ cells (Fig. 5C). Monocyte-derived myeloid-derived suppressor cells (MDSC) and TAMs have been demonstrated to inhibit the proliferation of T cells(*6*). Despite the alterations of cellular distributions within *Il1α^-/-^* tumors, a similar pattern of T cell proliferation suppression was observed (Fig. 5D-E). The Ly6C^+^MHC II^-^ quadrant exhibited the most potent suppressive activity on T cell proliferation, while Ly6C^-^MHC II^-^ cells displayed weaker suppressive effects, and Ly6C^+^MHC II^+^ and Ly6C^-^MHC II^+^ quadrants showed no apparent impact on T cell proliferation. It’s intriguing that co-culturing CD8 T cells with Mono/Mac cells prompted the upregulation of immune suppressive marker genes PD-1 and CTLA4 (Fig. 5F-G). However, the Ly6C^+^MHC II^+^ or Ly6C^-^MHC^+^ cells from tumors in *Il1α^-/-^*mice exhibited a diminished capacity to induce PD-1 and CTLA4 expression, indicating reduced immune suppressive abilities (Fig. 5D). Conversely, both Ly6C^+^MHC II^-^ and Ly6C^-^MHC II^-^ cells displayed minimal phagocytic ability, whereas Ly6C^+^MHC II^+^ or Ly6C^-^MHC^+^ cells demonstrated strong phagocytic capabilities. Interestingly, the Ly6C^+^MHC II^+^ cells from tumors in *Il1α^-/-^* mice showed a twofold decrease in phagocytosis compared to those from the wild type (Fig. 5H). This data aligns with prior research indicating that CX3CR1+ macrophages exhibit robust phagocytic capabilities in both normal mammary gland development and mammary tumors(*40, 41*). Collectively, these findings suggest that the absence of IL1α expression leads to the reprogramming of monocytes into immature macrophages with diminished phagocytic and immune suppressive capacities.

### Correlation with human monocytes differentiation and macrophage polarization

To further underscore the significance of our findings in human cells, we conducted the comparison analysis by selecting the top differentially expressed genes from the Mono, Mac1 through Mac6 clusters between the treatment groups and compared them with human myeloid cells subjected to various M1/M2 maturation triggers (Fig. 6A and Table S1). In total, 299 profiles from 32 different stimulation conditions were obtained (*14*). The enrichment analysis reveals that in the tumors from WT mice, clusters of Mono, Mac2, Mac4, Mac5, and Mac6 exhibited gene signatures, which were strongly associated with M2 maturation induced by IL10, IL13 and glucocorticoid (GC). In contrast, the corresponding clusters from *Il1α^-/-^* tumors displayed higher Z scores indicative of an inflammatory human monocyte signature triggered by cyclodextrin (MCD). Intriguingly, Mac3 cluster cells from tumors in both WT and *Il1α^-/-^* showed the strong enrichment for genes similar to human DC cells stimulated with TNFα and TNF/PGE2/Pam3CSK4 (TPP). Mac1cells from tumors in WT mice exhibited gene signatures resembling that of an unstimulated Monocyte and a variety of macrophages, while Mac 1 cells from Il1α ^-/-^ shared a gene signature with the activated human monocytes. Overall, these gene signatures effectively captured the essence of the less inhibitory immune phenotype of myeloid cells observed in the *Il1α^-/-^* TME, aligning closely with human myeloid cell signatures (Fig. 6B). To confirm these signature analysis, flow cytometry analysis was performed to examine the expression levels of M1 signature gene iNOS and M2 signature gene CX3CR1 in the tumor myeloid cells. Consistently we observed a significantly higher expression level of iNOS but a lower expression of CX3CR1 in *Il1α^-/-^* myeloid cells than WT cells (Fig. 6C-D). To determine whether the changes in the CX3CR1+ macrophage populations resulted from an altered ratio of classical (CX3CR1^hi^) and patrolling (CX3CR1^low/Neg^) monocytes from the blood, the expression levels in the respective myeloid cells from both tumor and blood were compared. While the CX3CR1+ myeloid population was reduced in *Il1α^-/-^* TME, no alterations were noticed in the cells from blood (Fig. S12). These results suggested that IL1α influences the differentiation of those recruited monocytes, shifting them away from the iNOS+ inflammatory phenotype and towards becoming CX3CR1+ immune suppressive macrophages(*41*). In line with our hypothesis that IL1α induces immune suppressive elevated expression of IL1α in primary cancers were indicative of an unfavorable prognosis for patients undergoing immunotherapy.

**Fig. 6.**
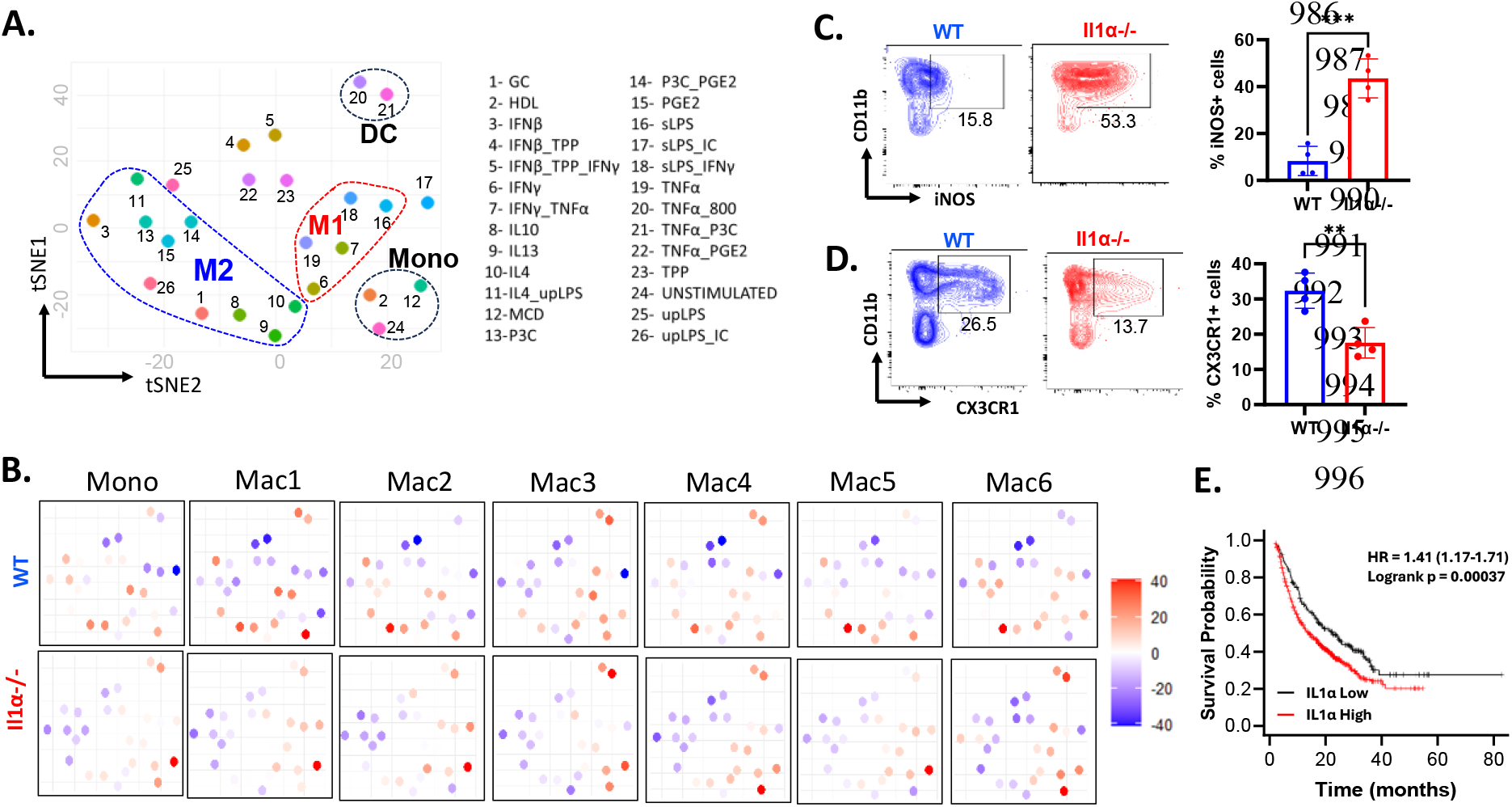
Il1α-mediated reprogramming of TAM phenotype and comparison to human macrophages. **A.** tSNE map juxtaposing dendritic cells and monocytes stimulated with diverse inflammatory triggers alongside human macrophages characterized by M1/M2 polarization gene profiles for GSE46903 dataset. **B**. tSNE map shown for each subcluster of mono/mac of WT and *Il1α^-/-^* in comparison to human macrophages under differing stimulation conditions. **C** and **D.** Percentage of CD11b^+^iNOS^+^ and CD11b^+^CX3CR1^+^ cells in tumor (n = 4). Unpaired Two Tailed t-test was performed. ***<0.05; **<0.05. **E**. Il1α expression levels predict prognosis of cancer patients after immunotherapy. Patients were stratified into two groups based on the mean values of Il1α expression (kmplot.com)(*53*).

## Discussion

In this study, we demonstrated that IL-1α derived from myeloid cells in the TME promotes breast tumor progression by regulating the immune response. Most previous studies have focused on the effects of tumor-derived IL-1α on malignant cells themselves or the proximal stroma(*2, 21, 22*). The contributions of host-derived IL-1α to tumorigenesis have been relatively underexplored. Early congenic graft experiments into WT and IL-1α knockout mice hinted the site-variable influences of host-derived IL-1α, with enhanced mammary oncogenesis yet suppressed melanoma progression(*27*). However, the underlying mechanisms on the pro-versus anti-neoplastic actions of host derived IL1α remain poorly defined. Our single cell RNA sequencing analyses clearly uncovered the sources of IL-1α within the TME of mammary tumors. We consistently detected abundant expression within CX3CR1^+^ macrophages infiltrating mammary tumors, rather than in the malignant cells themselves. Re-analysis of the previously published single cell RNA sequencing study of MMTV-neu tumors also showed IL-1α expression in CX3CR1^+^ macrophages and some neutrophils although these tumors were harvested at the different time points(*42*). However, neutrophils only account for a small fraction of infiltrating immune cells in our study. These data also align with single cell RNA sequencing studies in human breast cancer models highlighting myeloid cells as the foremost IL-1α reservoirs at tumor sites.

Given the dynamic changes in immune cell abundance and activity during cancer progression, we hypothesize that host-derived IL-1α assumes context-dependent functional roles at different stages of tumorigenesis. Using congenic engraftment models, we uncovered that impaired tumor growth in IL-1α knockout hosts stemmed from enhanced anti-tumor immunity. Single cell profiling and flow cytometry analysis consistently displayed increased immune infiltrates with greater abundance and activation of cytotoxic CD8 T cells within grafts in the absence of host IL-1α signaling. Moreover, selectively depleting CD4 and CD8 T cells, but not B cells, restored tumor expansion in IL-1α deficient animals. This implicates that adaptive cell-mediated responses are critical for immune control of tumors when host-derived IL-1α is lacking. Together, these findings demonstrate that host IL-1α dampens the endogenous anti-cancer immune response that is otherwise capable of rejecting implants.

The altered anti-tumor immunity is likely due to IL-1α-mediated reprogramming of mononuclear phagocytic cells in the TME. Single cell RNA sequencing analysis revealed a significant reduction in CX3CR1+ ductal-like macrophages (Mac6) and an increase in stress response-like macrophages (Mac2) in the absence of IL-1α. flow cytometry analysis further validated the change in myeloid cell subsets within the TME. This relative change in macrophage subsets is unlikely due to differential recruitment of CX3CR1^low^ classical and CX3CR1^high^ non-classical monocytes for the following reasons. First, the proportion of the CX3CR1-positive monocyte population is comparable in the blood of both wild type and IL-1α knockout mice. Secondly, IL-1α knockout mice showed mild leukopenia in both tumor-free and tumor-bearing conditions, consistent with the known role of IL-1α in hematopoiesis and mobilization of myeloid cells from the bone marrow (*31, 32*). In contrast, more immune cells infiltrate the tumors. Additionally, there are no significant differences or even a slight decrease in CCR2 expression in infiltrating monocytes or neutrophil of IL-1α knockout mice. Therefore, the change in infiltrating myeloid cell subsets is largely due to the results of local differentiation and/or proliferation of recruited monocytes or their precursors rather than differential recruitment.

Monocytes can differentiate into macrophages with a high degree of phenotypic and functional heterogeneity in both normal mammary gland and tumor tissues(*5, 41*). In normal mammary gland, there are two types of resident macrophages: a rare population of ductal (CX3CR1^+^CD11c^+^Ly6C^-^) and a major population of stromal (CX3CR1^-^CD11c^-^Ly6C^lo^MHC II^hi^) microphages(*38*). Interestingly, in response to change in their niches in tumors, only ductal macrophages drastically expanded while part of stromal macrophages (SM) change phenotype from CX3CR1^-^ to CX3CR1^+^ cells(*38*). We have observed a dramatic decrease in the DM-like population (Mac6) in the tumors of *Il1α^-/-^* mice, which is most likely due to selective inhibition of DM differentiation or proliferation. The increased stress response population (Mac2) in the *Il1α^-/-^* mice showed the stromal macrophage phenotype (CX3CR^-^Ly6C^lo^CD11c^lo/-^MHC II^hi/lo^). It is likely that knockout of *Il1α* leads to expansion of SM due to the altered niches(*40*). The enrichment of stress response and hypoxia pathway in the Mac2 population indicated that these cells are most likely exposed to hypoxia environment within the tumor. In contrast to the typical peripheral location of SM in the mammary tumors, we observed more macrophages inside the tumors from the *Il1α^-/-^* host mice.

The alterations in the differentiation routes of infiltrated monocyte may change their functions as well. Accumulating data suggests a continuous spectrum of functionally diverse macrophage states rather than ontogenically defined M1/M2 subsets in the TME(*9*). Most macrophages *in vivo* likely express markers of both M1 and M2 activation states(*16, 17*). CX3CR1 is a marker frequently associated with anti-inflammatory, immunosuppressive M2 macrophages and poor prognosis in breast cancer(*40, 43, 44*), whereas iNOS is a proinflammatory M1 macrophage biomarker that likely promotes anti-tumor immune responses(*45*). flow cytometry analysis showed that CX3CR1+ cells are intermixed with iNOS-positive cells although the number of CX3CR1+ and iNOS-expressing cells showed the opposite changes in the *Il1α^-/-^* mice in comparison with WT mice. In the expression profiles comparison with *in vitro*-induced human M1 and M2 macrophages, there is a general reduction in expression of M2 macrophage genes across the subpopulations although there are no distinct M1 and M2 subpopulations.

Monocyte-macrophage lineage of cells can suppress CTL responses through inhibition of proliferation and activation(*46, 47*). Several changes in monocyte-macrophage lineage of cells may explain the increased number and activation status of CTL in *Il1α^-/-^*mice. First, after sorting based on Ly6C and MHC-II expression, we observed the shift from the Ly6C^-^MHCII^-^ population to Ly6C^+^MHCII^+^ cells in the tumors from the *Il1α^-/-^* mice. Although the Ly6C^+^MHCII^+^ cells from both WT and *Il1α^-/-^* mice did not inhibit antigen-independent T cell proliferation, the Ly6C^-^MHCII^-^ subpopulations showed strong suppressive abilities on T cell proliferation. The relative cellular composition changes may partially explain the increased number of CD8 T cells in the transplants of *Il1α^-/-^* mice. Secondly, the CD8 T cells cocultured with Ly6C^+^MHC II^+^ population from WT mice expressed higher expression of exhaust markers such as PD-1 and CTLA4. Several previous reports indicate CX3CR1-positive TAMs commonly display high phagocytic but immune suppressive activities(*40, 48, 49*). In MMTV-PyMT models, monocyte derived (TAMs) present cancer cell antigens and drive exhaustion of cytotoxic T cell with induction of PD1 expression(*50*). Finally, *in vivo* depletion of TAMs, but not mammary tissue macrophages (MTMs), can efficiently restore the tumor infiltrating cytotoxic T-cell response by increasing the number of PD-1^-^Gzmb^+^ CD8 T cells(*37*). IL-1α deficiency-induced reprogramming of monocyte differentiation recapitulates some phenotypes of selective TAM depletion(*37*). Interestingly, ICI treatment can also remodel tumoral monocyte/macrophage lineage cells leading to selective ablation CX3CR1^+^CD206^+^ macrophages and accumulation of iNOS+ macrophages in an IFN-γ dependent manner(*15*). Similarly, our trajectory analysis revealed that loss of IL-1α skewed paths away from the CX3CR1+ MAC6 cluster toward the iNOS-positive Mac2 cluster, with Mac1 serving as a common progenitor enriched in IL-1 and IFNγ responsive signaling. This similarity raises the possibility that targeting IL-1α could be used to promote monocyte differentiation into proinflammatory macrophages or antigen-presenting myeloid cells to enhance the efficacy of immunotherapy.

Despite both scRNA seq and flow cytometry analysis showing a similar change trend, the relative proportions of CX3CR1 and iNOS-positive myeloid cells are much higher by flow cytometry compared to scRNA seq profiling. This difference may be due to some caveats of scRNA sequencing. First, mRNA and protein expression do not always directly correlate. We used relatively low cell count options for scRNA sequencing, which can cause large variations for genes with low abundance. Secondly, our sample digestion procedure caused substantial cell death. To improve single cell library quality, we used a live/dead staining kit to enrich live single cells. The tissue dissociation, live cell enrichment, and droplet encapsulation inherent to scRNA-seq caused under-representation of certain cell types including adipocytes, mast cells, tumor cells and some myeloid cell subsets(*51*). Given these systemic procedural biases, we therefore limited our analysis to relative changes between wild type and knockout grafts within the corresponding clusters, or to gene expression profiles within the same compartment. Furthermore, we validated the analysis using flow cytometry and verified it with published datasets from similar studies, extending the gene expression profiles of mouse monocytes/macrophages to their human counterparts. By comparing activation states of our macrophage clusters to published M1/M2 status markers of human equivalents, we obtained the results suggesting that the TME in IL-1α deficient mice likely polarizes macrophages toward an immune active phenotype. Further elucidating the mechanisms of IL-1α mediated immune suppression in the tumor microenvironment may lead to the development of new immunotherapeutic strategies for breast cancer.

## Materials and Methods

### Cell culture and mice

H605 MMTV-neu mammary tumor cell line in FVB/N background were cultured in DMEM/F12 media containing 10% fetal bovine serum, 10µg/mL insulin with 1% penicillin/streptomycin. *Il1α^-/-^* mice were kindly provided by Dr. Yoichiro Iwakura and have been backcrossed into FVB/N background as described previously(*21*).

### Breast cancer allograft mouse models

The animal studies were approved by the USC (University of South Carolina) committee for research in vertebrate animals. Cancer cells (2×10^6^ H605 cells) were mixed with Matrigel at 1:1 ratio and injected to the fourth mammary glands of WT or *Il1α^-/-^* FVB/N background mice. The tumor was measured every 3rd day and calculated using the formula-volume = ½(length x width^2^). The mice were sacrificed using cervical dislocation at the indicated time points. Blood, spleen and bone marrow were collected to study immune profile. For immune cells depletion, anti-CD19, CD4, and CD8 neutralization antibodies were administered at a dosage of 200µg per mouse. This treatment was initiated through intraperitoneal (i.p.) injection two days before tumor inoculation and continued at three-day intervals until day 25. Cellular depletions of CD8 T cells, CD4 T cells and CD19+ B cells were confirmed by flow cytometry of PBMC.

### Tissue collection and processing for immune profiling

Blood samples were collected into tubes coated with EDTA. For immune analysis, 100µL of blood was mixed with 900 µL of 1×RBC (red blood cell) lysis buffer. Upon quickly vortexing the sample at speed 5/6 on the vortex, the samples were placed on ice for 10 minutes and then centrifuge it at 500rcf for 5 minutes, leading to a white pellet at the bottom for flow cytometry analysis. For saving serum, first centrifuge the EDTA tube containing blood at 16,000g for 5 minutes. Carefully collect the supernatant into a clean tube, ensuring that the red blood cells remain undisturbed. Next, add 1ul of protein inhibitor prepared using Pierce^TM^ Protein Inhibitor tablets (A32955) for every 500µl of serum, and gently flick the tube to ensure thorough mixing. Store these samples at −80°C, taking care to avoid freeze-thaw cycles, to preserve the serum for further analysis.

The tumor processing protocol involved careful collection and division of the tumor into two equal parts. The first half was fixed in 4% Paraformaldehyde and left overnight at 4°C to prepare it for immune-fluorescence analysis. The second half was minced using a sterile razor in a 5cm dish. For cell isolation, the minced tumor was collected into a 15 ml tube and incubated in 5ml of digestion buffer, which contained 2mg/ml collagenase IV and 0.2mg/mL hyaluronidase in DMEM/F12, for 45 minutes at 37°C with agitation at 1000rpm in an Eppendorf shaker/incubator.

Following digestion, the tube was centrifuged at 500g for 5 minutes, and the supernatant was removed. The cell pellet underwent treatment with RBC lysis buffer, followed by resuspension in 5ml of wash buffer (consisting of 2% FBS in PBS) and filtration through a 40µm filter to obtain a single-cell suspension, which was then utilized for flow cytometry and cell sorting.

In the femur sample preparation process, both intact femurs were initially collected and cleaned by meticulously removing excess muscle and fat using a sterile razor. The cleaned femurs were grinded in a clean mortar in the presence of 5mL of wash buffer (2% FBS in 1× PBS). The resulting non-bone liquid phase from this grinding was collected and then filtered through a 40µm filter. The cell pellet obtained after centrifugation was resuspended in 1× RBC lysis buffer to eliminate red blood cells. The isolated cells were then counted and used for experiments.

The spleen was placed in a 5cm dish with 5ml of wash buffer. Clean non-charge glass slides were used to apply pressure on both sides to gently squeeze them into the wash buffer until the tissue becomes pale. The cells in wash buffer were subjected to centrifugation at 500g for 5 minutes, with careful removal of the supernatant. After RBC lysis, the remaining cells were resuspended in wash buffer and pass through 40µm filter to attain single cell suspension.

### Multiplex cytokine assay

Blood was collected into EDTA-treated tubes, plasma extracted, and samples frozen at −80°C until testing. Mouse cytokine and chemokine concentrations were determined using a multiplex immunoassay. 50 µl of plasma samples were sent for quantitative analysis using Eve Technologies Inc.

### Immuno-fluorescence analysis

The tumor samples were fixed in 4% PFA for 24 hrs and then switched to 30% sucrose for 30mins prior to O.C.T. embedding. Sections of 7 µm were cut using the cryosection microtome. Primary Antibodies, which are conjugated with a fluorophore (CD11b Cat No: 101206; 1:200, CD45 Cat No: 147708; 1:200) were used for immuno-fluorescents staining as described previously(*52*). A Zeiss LSM 800 confocal microscope equipped with 2.6ZEN software was used to acquire images.

### Flow cytometry

The single cell suspension from various tissues were quantified using Bio-Rad® automatic cell counter. A total of 1 million cells per stain were used, resuspended in 100µL of washing buffer for the staining of immune cells within the different tissues. A 1:1000 dilution of CD16/32 receptor blockade was applied and incubated for 5 minutes, followed by the addition of the desired antibody at the appropriate dilution to the single cell suspension (**Table S2**). The cells were then incubated for 15 minutes at 4°C and subsequently centrifuged at 500g for 5 minutes. After that, a live-dead stain (Zombie Aqua/Yellow/7-AAD) at 1:1000 dilution was used to incubate with cells for 10 minutes at room temperature in the dark. The cells were centrifuged again at the same speed and duration. To assess T cell cytokine production capability, activation cocktail and golgi plug were incubated with cells for 1 hour as per the manufacturer’s instructions. Finally, the samples were fixed using BD Cytofix/Cytoperm Plus (#555028) for intracellular protein staining, following the manufacturer’s guidelines. Nuclear proteins were analyzed using the eBioscience™ Foxp3 intracellular stain kit (#00-5523-00), following the manufacturer’s instructions (**Table S3**).

### scRNA seq and analysis

Single cells were extracted from two sets of tumor samples derived from both WT and *Il1α^-/-^* mice, collected at 14 days post-transplantation. Elimination of dead cells was carried out using the dead cell removal kit (#130-090-101, Miltenyi Biotec), following the manufacturer’s guidelines. Cells with viability greater than 90% were used and kept on ice for fixation and single cell RNA-Seq analysis. Droplet-based single-cell partitioning and single-cell RNA-Seq libraries were generated using the Chromium Single-Cell 3′ Reagent v2 Kit (#PN-1000269, 10× Genomics, Pleasanton, CA) per the manufacturer’s protocol. The size profiles of the pre-amplified cDNA and sequencing libraries were examined by Agilent Bioanalyzer 2100 using a High Sensitivity DNA chip (Agilent). Samples were sequenced up to mean read of 23k reads per sample.

The sequencing data was analyzed using the Cell Ranger Pipeline (version 2.0.1) to perform quality control, sample demultiplexing, barcode processing, alignment, and single-cell 3′ gene counting. All sequencing data processing steps were conducted with Seurat v4.3. Initially, quality control was carried out on each library to establish appropriate filtering criteria. Expression matrices for individual samples were imported into R as Seurat objects, retaining only cells with a gene count of more than 200. Cells with poor quality, characterized by more than 5% mitochondrial genes, were excluded. Genes that were not expressed in at least three cells were also removed. To mitigate the technical variation, sequencing depth, and capture efficiency biases, we employed NormalizeData for normalization, and scaling according to Seurat protocol. For distinction, we assigned identifiers for WT and *Il1α^-/-^* samples. After quality control and integration, we analyzed a total of 13,335 cells. Within the tumor, 14 clusters were identified. Visualization of cell clustering was achieved using uniform manifold approximation and projection (UMAP). To assign cluster identities, we initially compiled a list of established cell types, along with their known markers. We assessed the expression of these markers, as well as additional canonical markers, using the FindAllMarkers() function in Seurat. Differential expression of each cluster was compared between WT and *Il1α^-/-^* cells by using log1p(AverageExpression()). GO analysis was performed using unique genes from FindAllMarkers() function in subcluster Mono/Mac using ClusterProfiler library and enrichGO function. The raw values 75from GO analysis was transported into software.broadinstitute.org/morpheus/ online software to make heat map. Representative genes graphs were made using violnplot(), DoHeatmap() and dotplot() function.

Monocle3 package was used on Seurat object. Myeloid subset object was treated as cell data set for each sample using function as.cell_data_set function and cluster_cells was used to make UMAP for the converted cell data set. The learn_graph function was used to predict the trajectory pathway. Used function order_cells to select Mono population as starting of trajectory and graph the trajectory as result.

### *In vitro* T cell proliferation assay

Miltenyi bead separation was used according to the manufacturer’s instructions to purify CD8 T cells from the spleen tissue of WT FVB mice (#130-104-075, Miltenyi Biotec). Four subsets of tumor-infiltrated myeloid cells (CD45^+^CD11b^+^Ly6G^-^) were sorted out based on the expression levels of Ly6C and MHC II. 1×10^6^ of CD8 T cells were labeled with 500 μL of 5μM carboxyfluorescein diacetate succinimidyl ester (CFSE; eBioscience). The CFSE-labeled T cells were then seeded at 1:4 ratio (T cells: Myeloid) in a non-adherent 96-well plate. Stimulation of these cells was achieved by adding CD3+CD28 MACSibead particles (#130-093-627) according to manufactures protocol. After three days of culture, the cells were subjected to centrifugation and stained for CD8 and exhaustion/activation markers, alongside a live/dead marker such as DAPI or Live/dead stain, to assess T-cell proliferation among live cells. Percentage suppression of proliferation with myeloid cells is calculated as (1-Proliferation with myeloid cells/Proliferation without myeloid cells)×100.

### Phagocytosis Assay

Myeloid cell subsets, isolated from tumor infiltration and distinguished by their expression levels of Ly6C and MHC II, were plated and treated with 25μm sized GFP-encapsulated particles at a ratio of 10 particles per cell in 100µL of media in a non-adherent 96 well plate. The cells were incubated at 37°C for 2 hours and flow cytometry was conducted on CD11b stained versus GFP to assess phagocytosis.

### Human macrophage M1/M2 gene signature analysis

Human expression data was accessed from GEO with GSE46903 ID. The samples from the time point of 72 hours treated with GM-CSF were used for M1/M2 macrophage status along with dendritic cells and monocytes with different triggers to construct the tSNE for analysis. The top 100 differential expressed genes from Mono/Mac subclusters between *WT* and *Il1α^-/-^* were used to calculate the z-score for each cluster with respect to the sample. The plots were made using the ggplot2 package on the constructed tSNE plot.

### Statistical analysis

All quantitative data are presented as mean± SD or SEM as indicated. Prism v10 was used to perform 2-way ANOVA and unpaired two tailed t-test for all graphs used. Survival curves were evaluated using the Kaplan–Meier method, and the differences between those survival curves were tested by the log-rank test.

## Acknowledgments

We are deeply grateful for the technical assistance provided by Drs. Michael Shtutman and Diego Altomare in single-cell sequencing, as well as Dr. Jason Kubinak in flow cytometry analysis.

## Funding

National Institutes of Health grant R01CA266027 (IR, EB, CH) National Institutes of Health grant R21 CA252360 (CH, PX)

## Author contributions

Conceptualization: MK, GZ, HC

Methodology: MK, GG, GA, KK, AA, MG, IR, EB, MC, HJ, CL, HW, DF, PX, JL, CH

Investigation: MK, GG, GZ, CH

Visualization: MK, CH Supervision: CH Writing—original draft: MK, CH

Writing—review & editing: HW, DF, GZ, CH

## Competing interests

Authors declare that they have no competing interests.

## Data and materials availability

The accession number for the scRNAseq data reported in this paper is GEO: GSE 264177.

## Supplementary Materials for

**Fig. S1.**
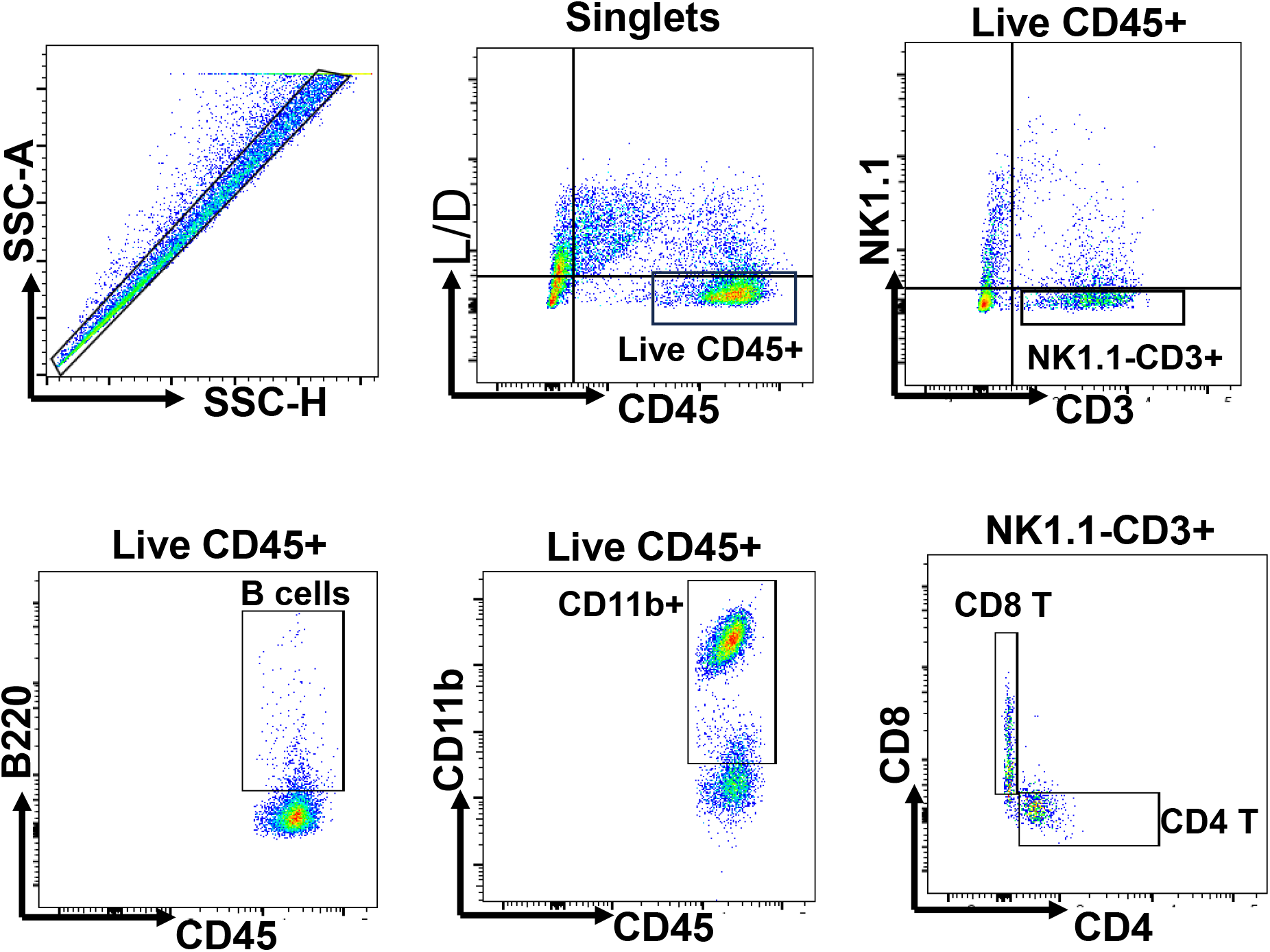
Tumor Immune cells gating strategy. Tumor single cells stained from 2-week time point to study overall immune lineage such as B cells (B220+), CD4 and CD8 T cells, Myeloid cells (CD11b+) which are singlets and Immune cell marker positive (CD45+) live cells.

**Fig. S2.**
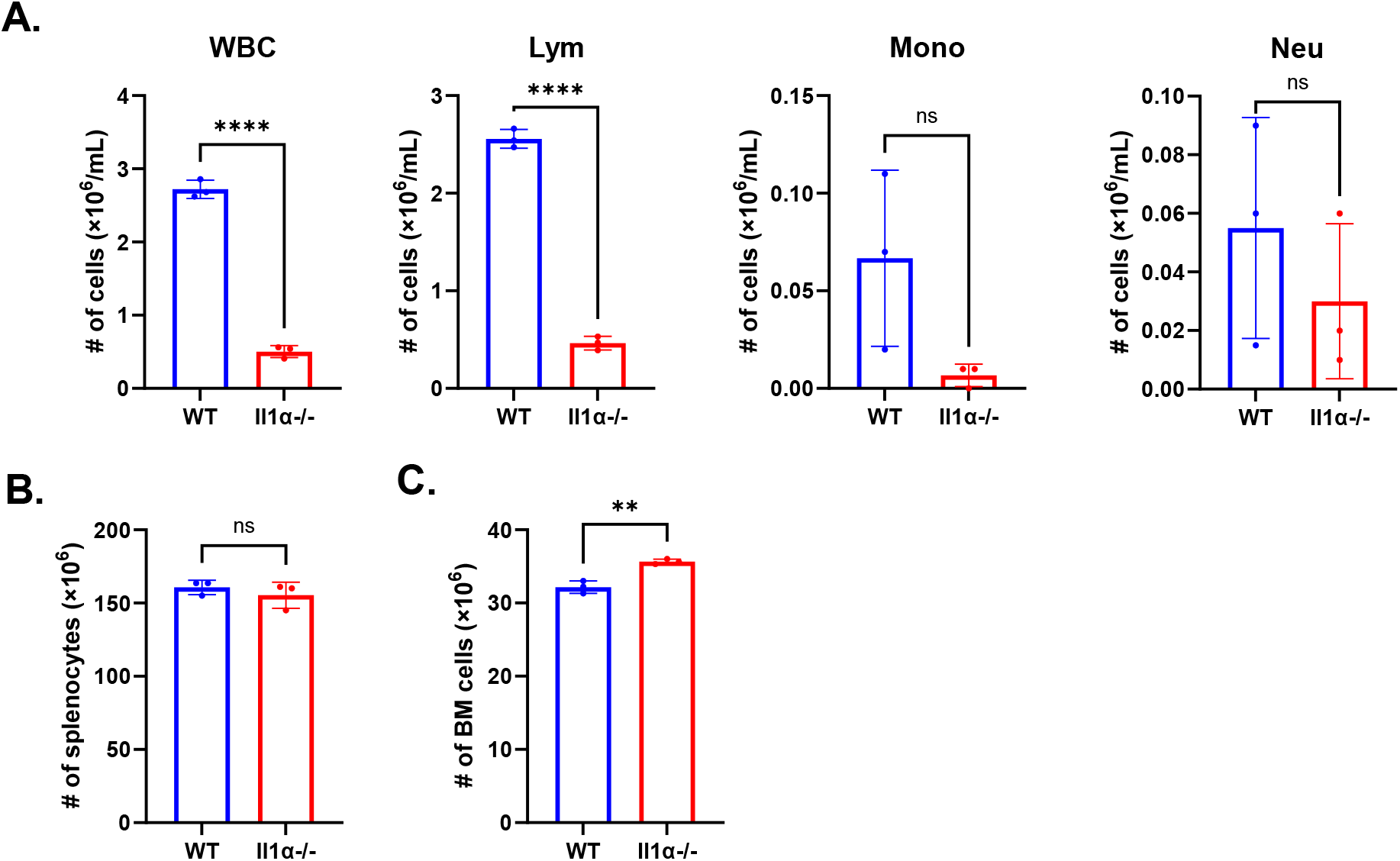
Immune profiles of tumor-free WT and Il1α^-/-^ mice. **A.** Blood Vetscan® data shown between tumor-free WT and *Il1α^-/-^* mice. **B.** and **C** describe the total number of spleen and bone marrow leukocytes from tumor-bearing WT and *Il1α^-/-^* mice. Unpaired Two Tailed t-test was performed. **** p<0.0001; **p < 0.01; ns, not significant.

**Fig. S3.**
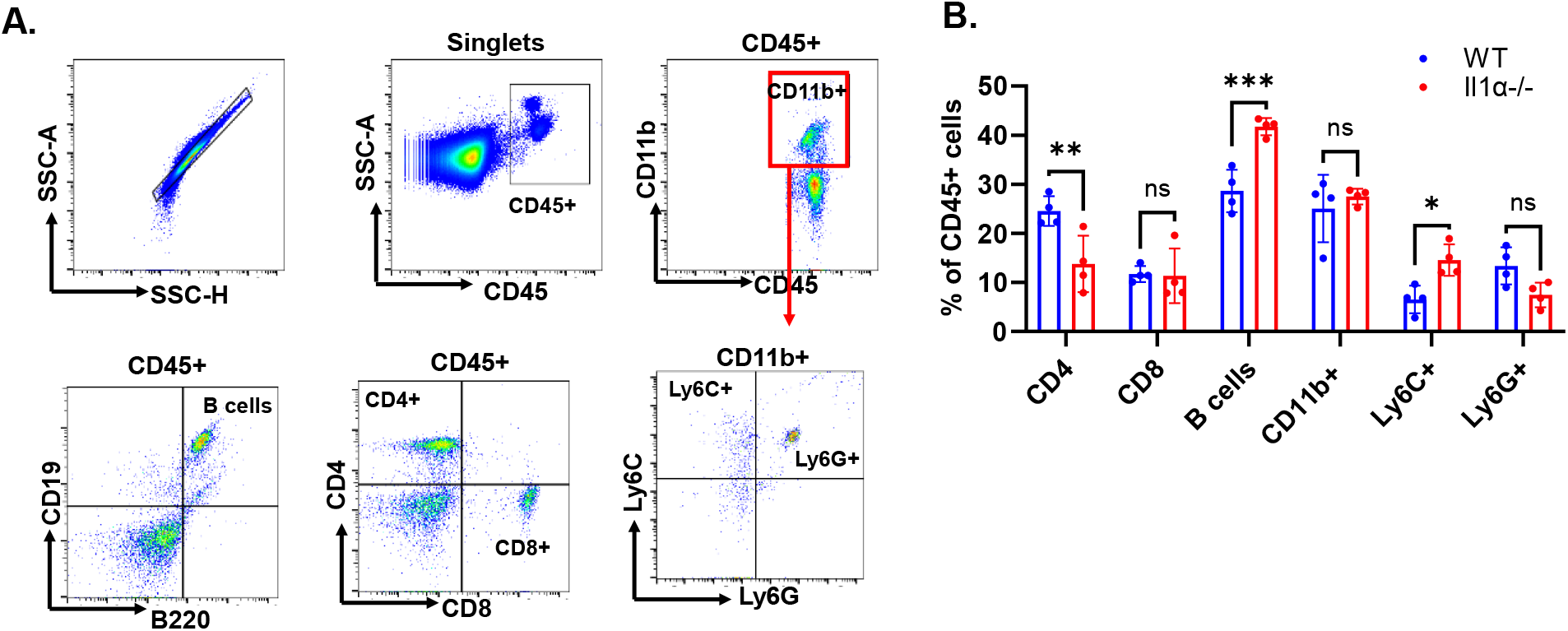
Blood Immune cells gating strategy. **A.** Blood collected from 2-week time point to study overall immune lineage such as B cells (B200+), CD4 and CD8 T cells, Myeloid cells (CD11b+) and their subsets Ly6C+ and Ly6G+ which are singlets and Immune cell marker positive (CD45+) live cells **B**. Quantified distribution of immune cells in blood comparing *WT* and *Il1α^-/-^.* Unpaired Two Tailed t-test was performed. ***<0.001; **<0.01.

**Fig. S4.**
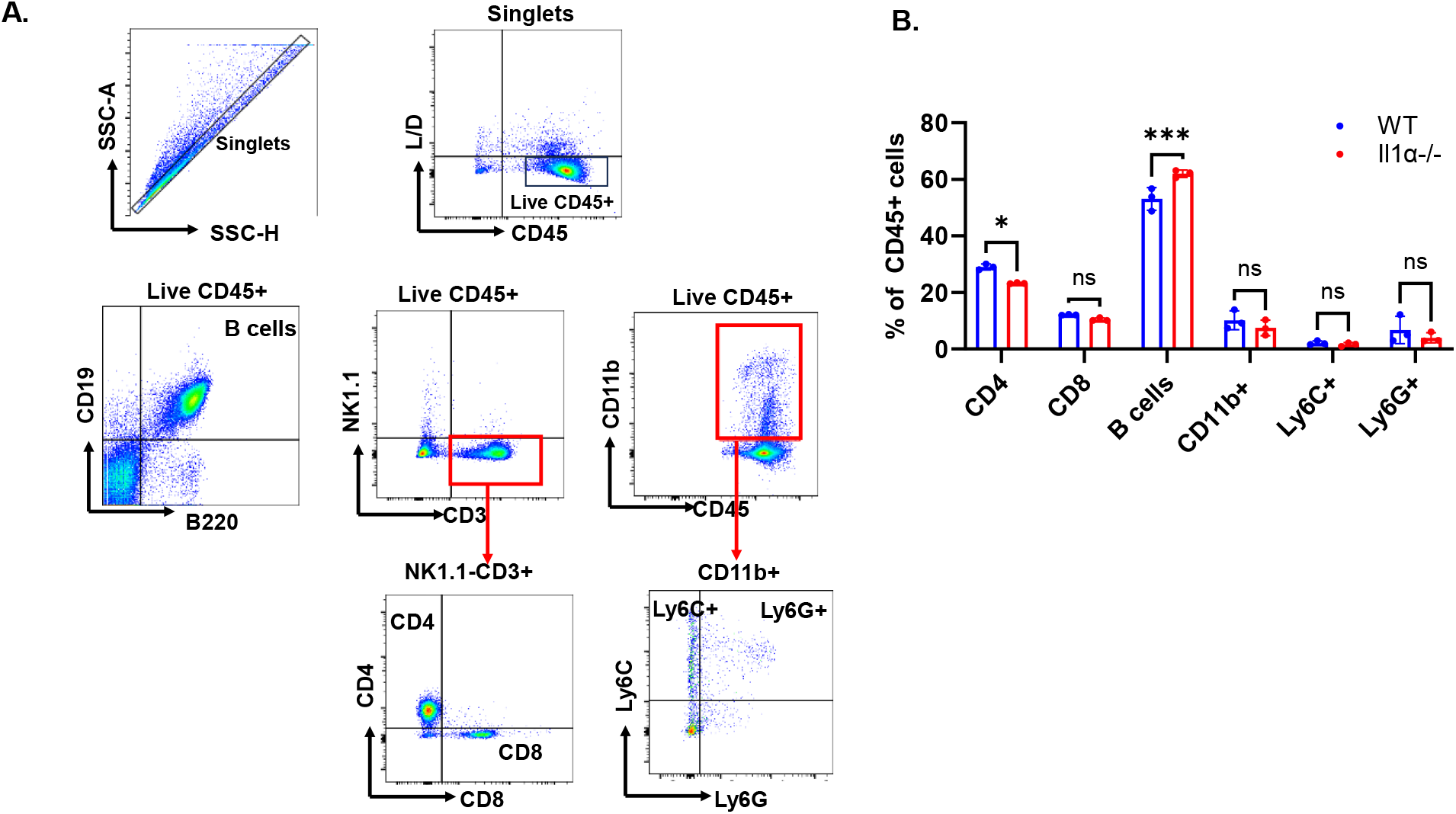
Spleen Immune cells gating strategy. **A.** Spleen from 2-week time point to study overall immune lineage such as B cells (B200+), CD4 and CD8 T cells, Myeloid cells (CD11b+) and their subsets Ly6C+ and Ly6G+ which are singlets and Immune cell marker positive (CD45+) live cells **B**. Quantified distribution of immune cells in spleen comparing *WT* and *Il1α-/-*. Unpaired Two Tailed t-test was performed. ***<0.001; **<0.01.

**Fig. S5.**
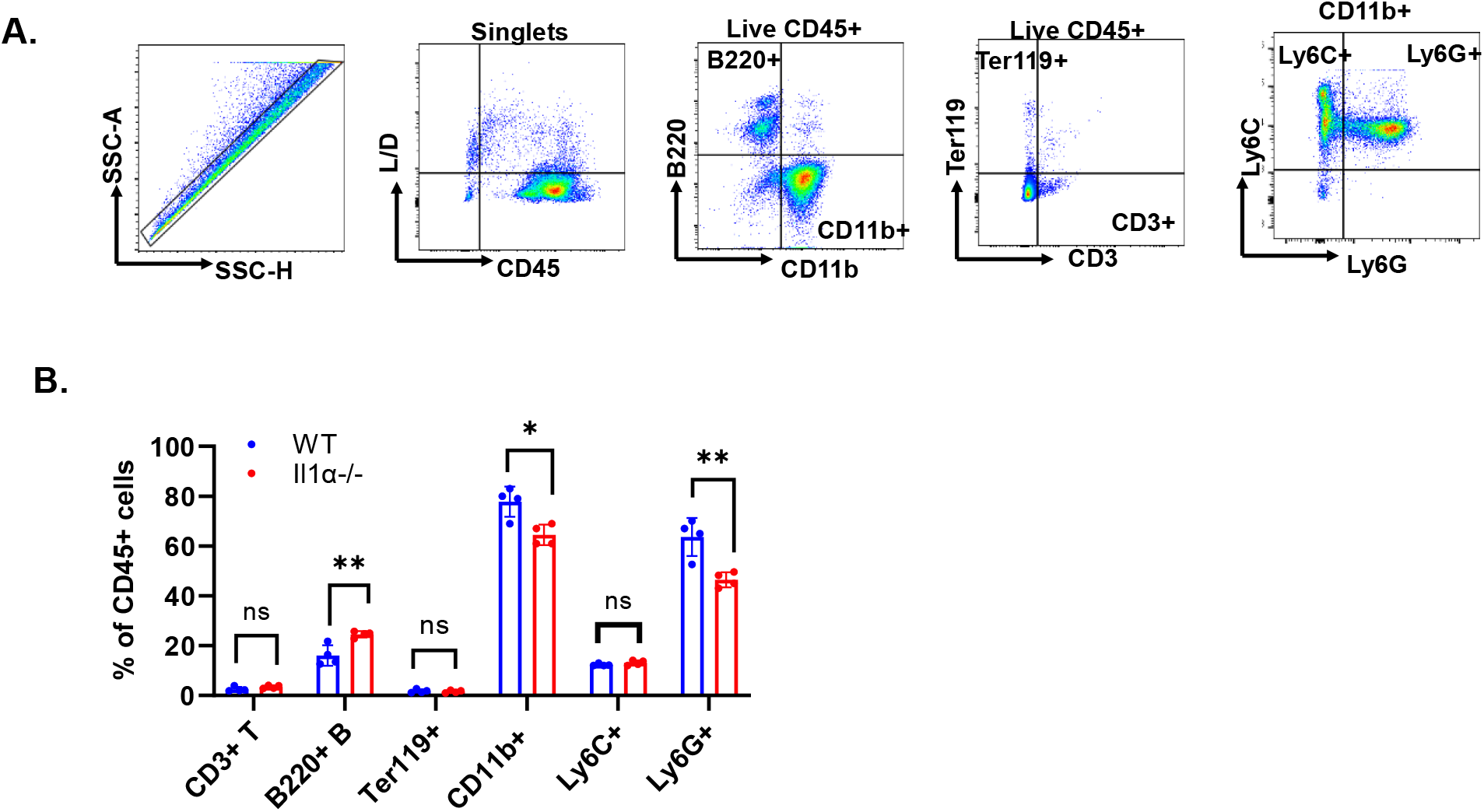
Bone Marrow Immune cells gating strategy. **A.** Bone marrow from 2-week time point to study overall immune lineage such as B cells (B200+), CD3+ T cells, Myeloid cells (CD11b+), Erythrocytes (Ter119+) and subsets of myeloid cells Ly6C+ and Ly6G+ which are singlets and Immune cell marker positive (CD45+) live cells. **B**. Quantified distribution of immune cells in bone marrow comparing *WT* and *Il1α^-/-^* .Unpaired Two Tailed t-test was performed. ***<0.001; **<0.01.

**Fig. S6.**
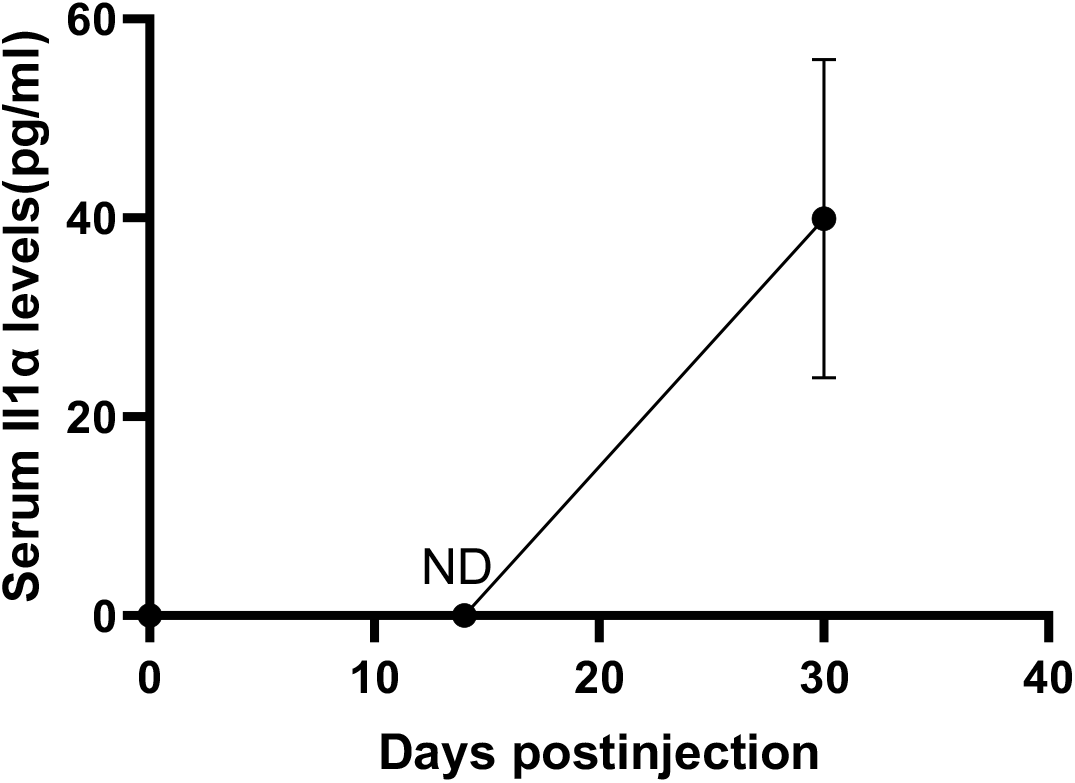
Il1α plasma dynamics. Plasma was monitored from day 0 until day 30 from *WT* tumor bearing mice quantified using Multiplex ELISA assay for Il1α (n = 3). ND, not detectable using this technology. The expression levels are not notably elevated compared to the background.

**Fig. S7.**
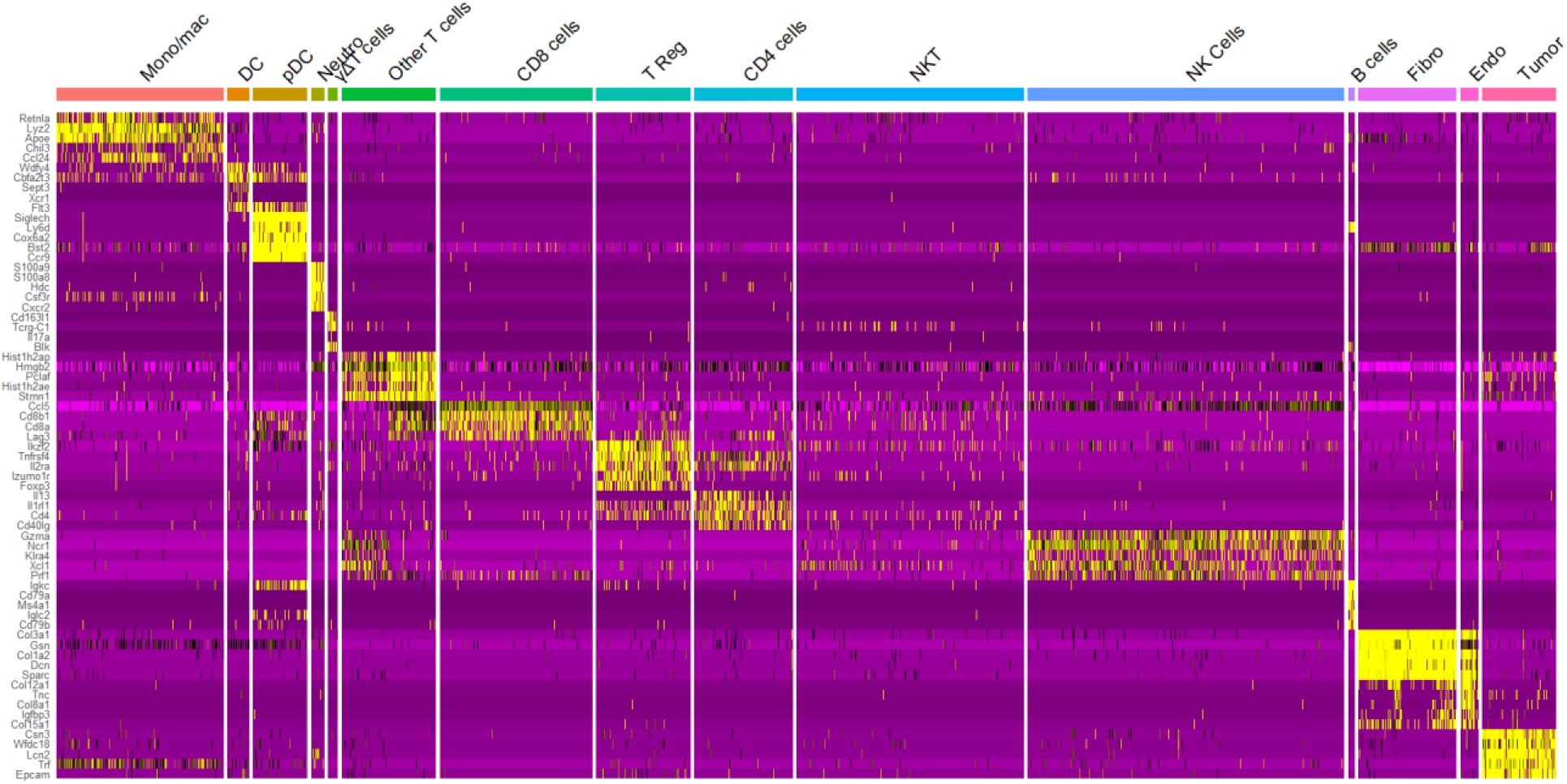
The top unique genes from the 14 overall clusters shown in heat map from scRNA seq analysis performed on 2-week tumor.

**Fig. S8.**
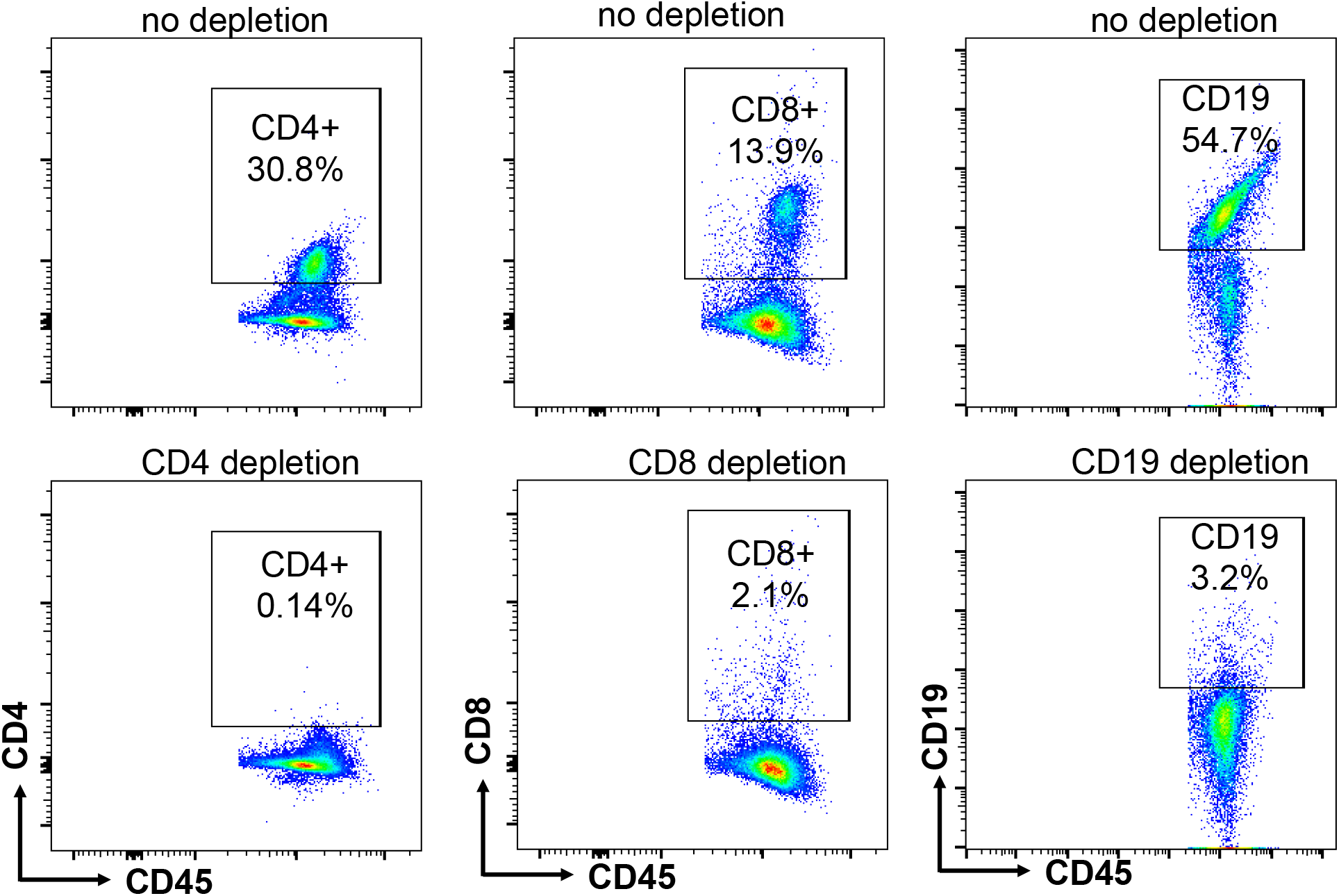
Immune cell depletion validation. Representative flow cytometry data of spleen to indicate depletion of CD4, CD8 and CD19 using neutralization antibody pre-gated for CD45+ live cells.

**Fig. S9.**
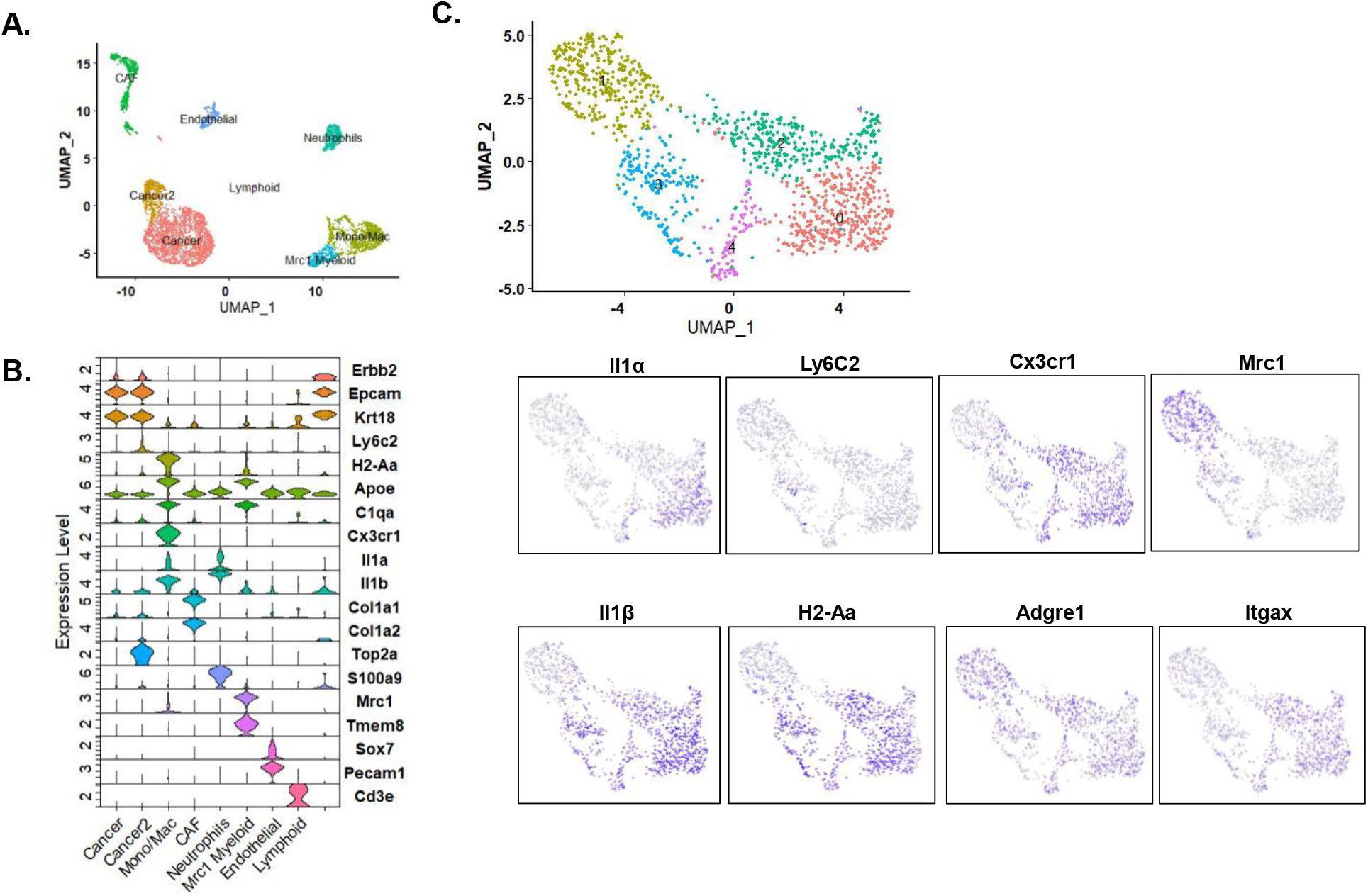
Cross verified MMTV driven HER2+ mice tumor scRNA seq dataset for Il1α source from GEO accession number: GSE166321. **A.** The representative tSNE, **B.** The top genes are shown in violin plot to identify the clusters. **C.** The Mono/Mac and Mrc1+ myeloid clusters were reclustered to find 5 subclusters as shown in the UMAP with genes Il1a, Ly6c2, Cx3cr1, Mrc1, Il1b, H2-Aa, Adgre1 and Itgax.

**Fig. S10.**
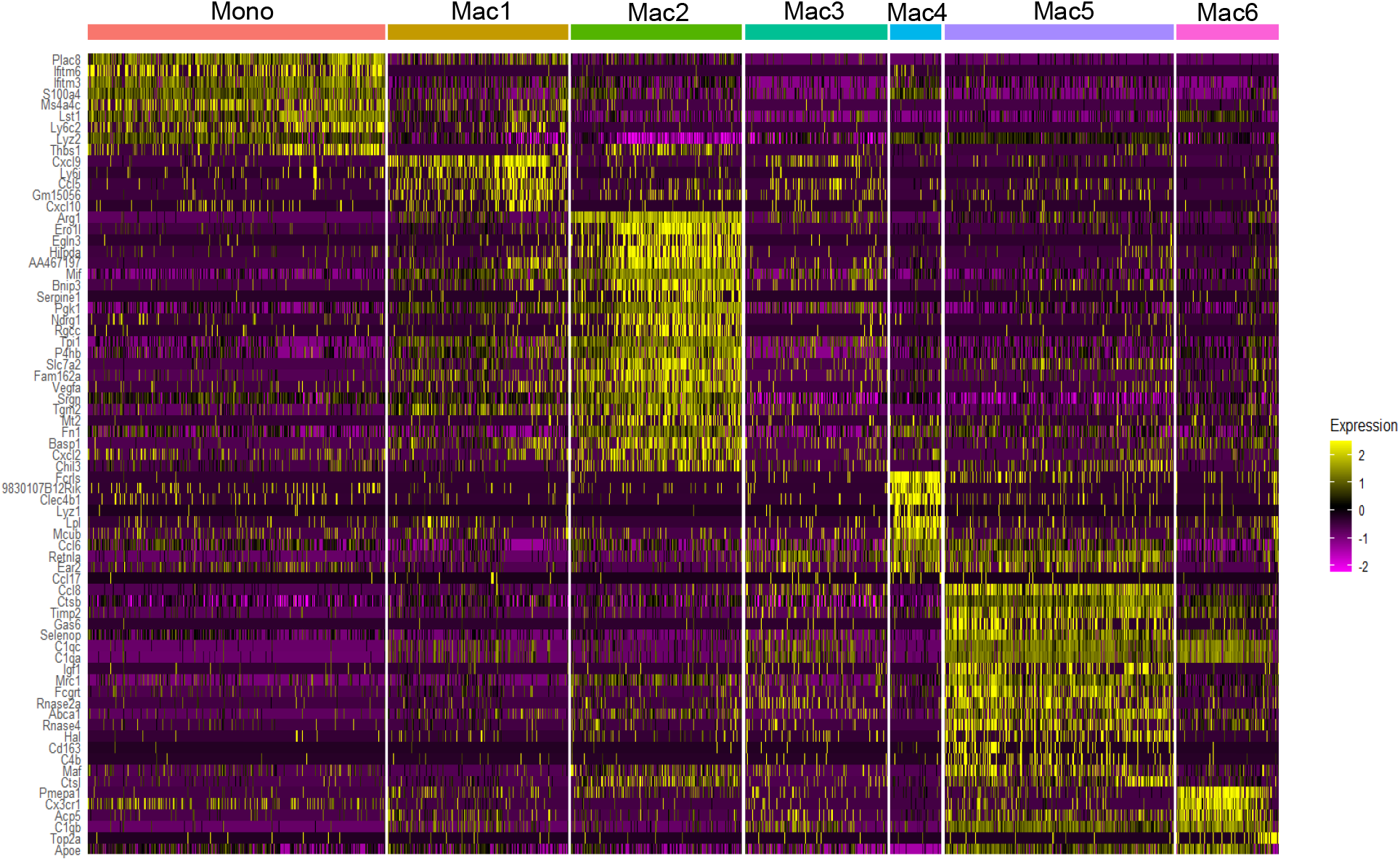
Top unique genes from the reclustered mono/mac subclusters from the 2-week time point tumor shown as heat map using scRNA seq analysis.

**Fig. S11.**
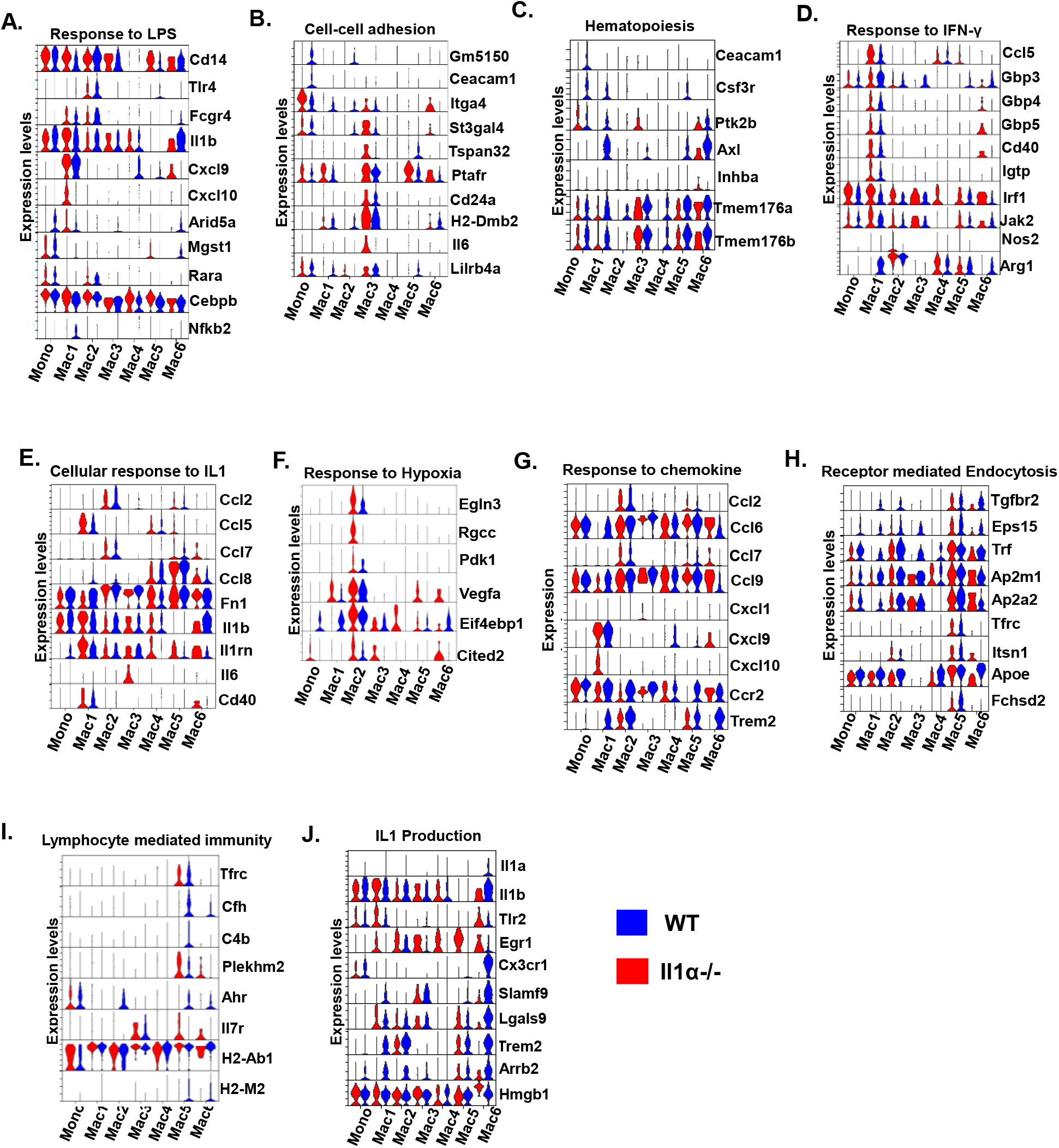
Unique genes selected to show from GO-pathway analysis with respect to WT (Blue) and *Il1α^-/-^* (Red) in Violin plot from scRNA seq analysis.

**Fig. S12.**
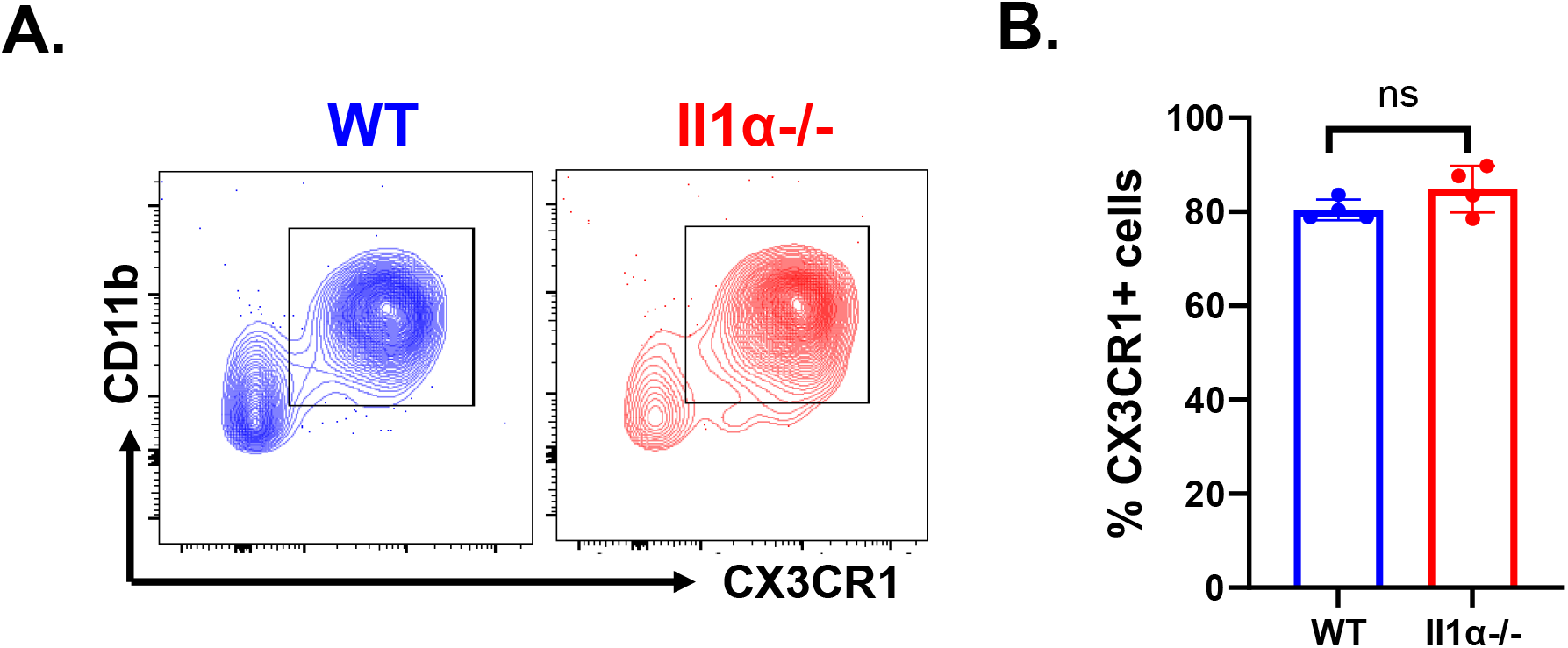
CX3CR1+ myeloid cells in blood from 2-week time point are not quantitively different between the groups. **A.** Representative figure of CD11b+Cx3cr1+ flow cytometry from blood which is pre-gated from CD45+ live cells. **B.** Quantified percent of CD11b+Cx3cr1+ cells in blood n = 4. Unpaired Two Tailed t-test was performed.

---

**Table S1.**
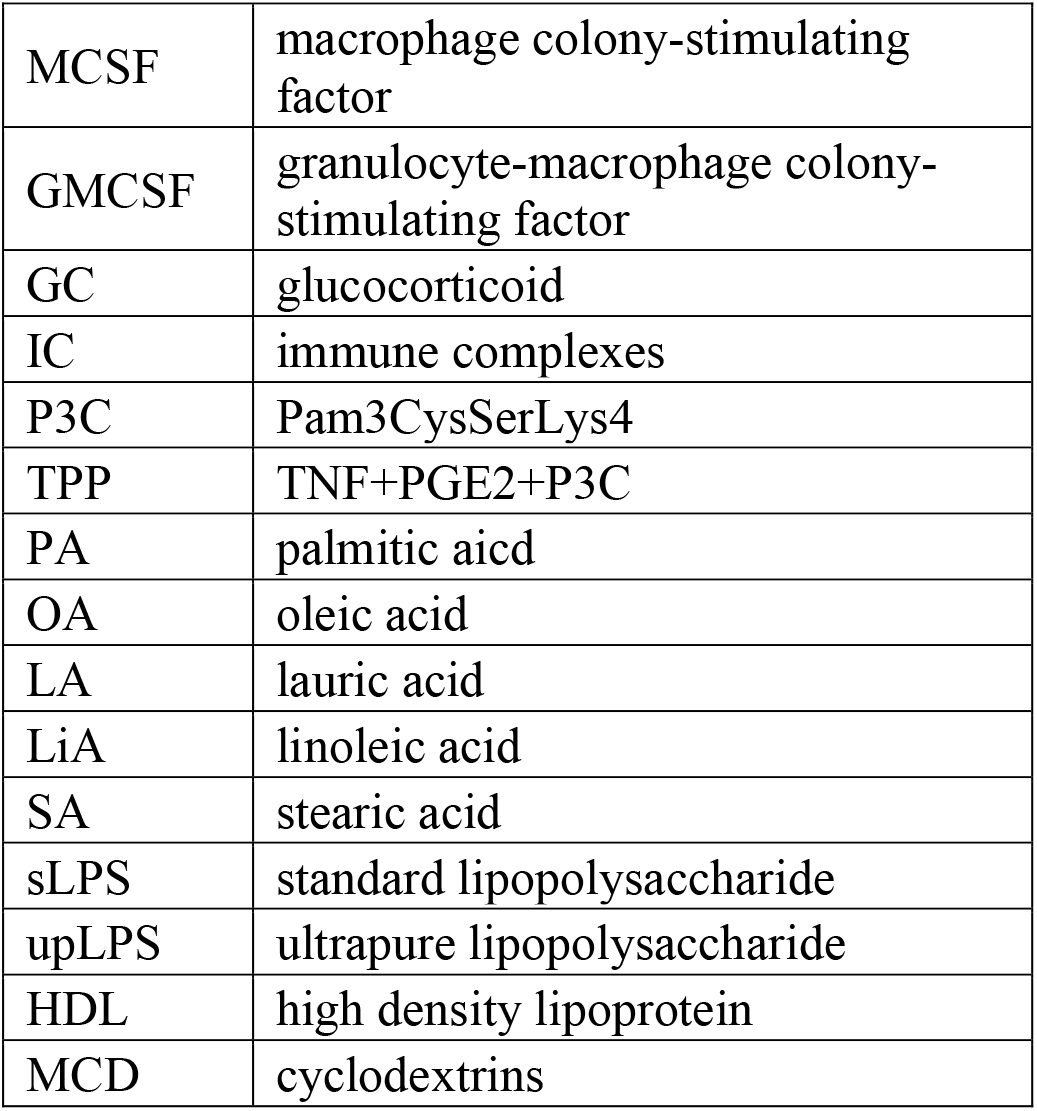
Abbreviations for stimuli used for induction of human macrophage polarization.

|  |  |
| --- | --- |
| MCSF | macrophage colony-stimulating factor |
| GMCSF | granulocyte-macrophage colony-stimulating factor |
| GC | glucocorticoid |
| IC | immune complexes |
| P3C | Pam3CysSerLys4 |
| TPP | TNF+PGE2+P3C |
| PA | palmitic acid |
| OA | oleic acid |
| LA | lauric acid |
| LiA | linoleic acid |
| SA | stearic acid |
| sLPS | standard lipopolysaccharide |
| upLPS | ultrapure lipopolysaccharide |
| HDL | high density lipoprotein |
| MCD | cyclodextrins |

**Table S2.** List of antibodies used in the study.

| **Target** | **Host** | **Clone** | **Fluor** | **Catalog Number** | **Company** | **Final Dilution** |
| --- | --- | --- | --- | --- | --- | --- |
| Arg1 | Rat | AlexF5 | PE | 12-3697-80 | Invitrogen | 1:100 |
| B220 | Rat | RA3-6B2 | FITC | 103206 | Biolegend | 1:200 |
| CD11b | Rat | M1/70 | FITC | 101206 | Biolegend | 1:400 |
| CD11b | Rat | M1/70 | APC | 101212 | Biolegend | 1:400 |
| CD11b | Rat | M1/70 | PE | 101208 | Biolegend | 1:400 |
| CD11c | Armenian Hamster | N418 | PE/Cy7 | 117318 | Biolegend | 1:100 |
| CD11c | Armenian Hamster | N418 | BV421 | 117330 | Biolegend | 1:100 |
| CD19 | Rat | 6D5 | PE/Cy7 | 115520 | Biolegend | 1:100 |
| CD3 | Armenian Hamster | 145-2C11 | PE | 100308 | Biolegend | 1:400 |
| CD3 | Rat | 17A2 | FITC | 100204 | Biolegend | 1:400 |
| CD4 | Rat | RM4-5 | FITC | 100510 | Biolegend | 1:100 |
| CD40L | Rat | SA047C3 | PE/Cy7 | 157008 | Biolegend | 1:100 |
| CD44 | Rat | IM7 | BV510 | 103044 | Biolegend | 1:100 |
| CD45 | Rat | 30-F11 | APC | 103112 | Biolegend | 1:400 |
| CD8a | Rat | 53-6.7 | APC/Cy7 | 100714 | Biolegend | 1:100 |
| CTLA4/CD152 | Armenian Hamster | UC10-4B9 | BV421 | 106312 | Biolegend | 1:100 |
| CX3CR1 | Mouse | SA011F11 | BV421 | 149023 | Biolegend | 1:100 |
| CX3CR1 | Mouse | SA011F11 | PE/Cy7 | 149016 | Biolegend | 1:100 |
| F4/80 | Rat | BM8 | APC/Cy7 | 123118 | Biolegend | 1:100 |
| F4/80 | Rat | BM8 | BV510 | 123135 | Biolegend | 1:100 |
| F4/80 | Rat | BM8 | PE | 123110 | Biolegend | 1:100 |
| GR1 | Rat | RB6-8C5 | PE | 108408 | Biolegend | 1:400 |
| Granzyme B | Mouse | OA16A02 | PE/Cy7 | 372214 | Biolegend | 1:100 |
| IFNγ | Rat | XMG1.2 | BV421 | 505830 | Biolegend | 1:100 |
| IL1a | Armenian Hamster | ALF-161 | PE | 503203 | Biolegend | 1:100 |
| IL1b | Rat | NJTEN3 | PE | 12-7114-80 | Invitrogen | 1:100 |
| iNOS | Rat | W16030C | PE | 696806 | Biolegend | 1:100 |
| KI67 | Rat | 16A8 | BV421 | 652411 | Biolegend | 1:100 |
| Ly6C | Rat | HK1.4 | BV785 | 128041 | Biolegend | 1:100 |
| Ly6C | Rat | HK1.4 | PE/Cy7 | 128018 | Biolegend | 1:100 |
| LY6G | Rat | 1A8 | APC/Cy7 | 127624 | Biolegend | 1:100 |
| LY6G | Rat | 1A8 | BV510 | 127633 | Biolegend | 1:100 |
| LY6G | Rat | 1A8 | BUV387 | 127678 | Biolegend | 1:100 |
| MHCII | Rat | M5/114.15.<br>2 | PE/Cy7 | 107630 | Biolegend | 1:200 |
| MHCII | Rat | M5/114.15.<br>2 | FITC | 107606 | Biolegend | 1:200 |
| NK1.1 | Mouse | S17016D | PE/Cy7 | 156514 | Biolegend | 1:100 |
| PD-1 | Rat | RMP1-30 | PerCP/Cy5.<br>5 | 109120 | Biolegend | 1:100 |
| Ter119 | Rat | TER-119 | PE/Cy7 | 116221 | Biolegend | 1:100 |
| CD4 | Rat |  | GK1.5 | BE0003-1 | Bioxcell |  |
| CD8a | Rat |  | 2.43 | BE0061 | Bioxcell |  |
| CD19 | Mouse |  | 4G7 | BE0281 | Bioxcell |  |

**Table S3.** List of key reagents used in this study.

| **Reagent** | **Supplier** | **Catalog Number** | **Final Dilution** |
| --- | --- | --- | --- |
| DMEM/F12 | Corning | 10-090-CV |  |
| RPMI | Corning | 10-040-CV |  |
| Fetal Bovine Serum | Peak Bio | PS-FB3 |  |
| Recombinant Human Insulin | Gibco | Rp-10908 |  |
| Phosphate Buffered Saline | Corning | 21040CV |  |
| Penicillin/Streptomycin Solution | Corning | 30-002-CI |  |
| Antibiotic Antimycotic Solution | Corning | 30-004-CI |  |
| 0.25% Trypsin | Gibco | 25200-072 |  |
| Matrigel Matrix | Corning | CB-40234 |  |
| 1ml 25G syringe | BD | 309626 |  |
| 0.5ml 28G syringe | BD | 329461 |  |
| EDTA | VWR | E522-100ML |  |
| Pierce™ Protein Inhibitor tablets | Thermo Scientific | A32955 |  |
| EDTA Blood Collection Tubes | BD | BD367842 |  |
| Vetscan HM5 Hematology Analyzer | Abaxis | 770-0000 |  |
| 10X RBC lysis buffer | Biolegend | 420302 |  |
| 4% Paraformaldehyde in PBS | Thermo Scientific | AAJ61899AK |  |
| No.9 Razor Blades | Garvey | 40475 |  |
| 5cm culture dish | Corning | 430166 |  |
| 15ml tubes | Extra Gene | P1013-15BF |  |
| Collagenase IV | Worthington Biochem | LS004188 |  |
| Hyaluronidase | MP Biochemicals | 100740 |  |
| 40Micron Filter | VWR | 21008-949 |  |
| Sucrose | Macron | MK772304 |  |
| Peel-A-Way OCT Mold | Polysciences | 18646A |  |
| Tissue-Tek OCT compound | Sakura | 4583 |  |
| Cryosection station | Leica | CM1860UV |  |
| ColorMark Charged Glass Slides | Epredia | CM-4951WPLUS-001 |  |
| Uncharged Glass Slides | Fishcer Scientific | 12-544-4 |  |
| Cell Counter | Bio-Rad | TC20 |  |
| Flow cytometry Machine | BD | FACSAria II |  |
| Flow cytometry Machine | BD | Symphony A5 |  |
| Cell sorter | Sony | SH800 |  |
| Zombie Aqua | Invitrogen | L34957 | 1:1000 |
| Zombie Yellow | Invitrogen | L34968 | 1:1000 |
| 7-AAD | Biolegend | 420404 | 1:1000 |
| Cytofix/cytoperm plus kit | BD | 555028 |  |
| Foxp3 intracellular stain kit | eBioscience | 00-5523-00 |  |
| cd16/32 receptor blockade | BD | 553142 | 1:1000 |
| CD8 isolation kit | Miltenyi Biotec | 130-104-075 |  |
| carboxyfluorescein diacetate succinimidyl ester (CFSE) | eBioscience | 65-0850-84 |  |
| Anti-CD3/CD28 beads | Miltenyi Biotec | 130-093-627 |  |

## Notes

### Competing Interest Statement

The authors have declared no competing interest.

